# Perturbation of the gut microbiota by antibiotics results in accelerated breast tumour growth and metabolic dysregulation

**DOI:** 10.1101/553602

**Authors:** Benjamin M Kirkup, Alastair McKee, Kate A Makin, Jack Paveley, Shabhonam Caim, Cristina Alcon-Giner, Charlotte Leclaire, Matthew Dalby, Gwenaelle Le Gall, Anna Andrusaite, Peter Kreuzaler, Avinash Ghanate, Paul Driscoll, James MacRae, Enrica Calvani, Simon WF Milling, Mariia Yuneva, Katherine N Weilbaecher, Tamas Korcsmáros, Lindsay J Hall, Stephen D Robinson

## Abstract

**Background:** Breast cancer is the second most prevalent cancer worldwide with around 1.7 million new cases diagnosed every year. Whilst prognosis is generally favourable in early stages, this worsens significantly in advanced disease. Therefore, it is pertinent to focus on mitigating factors that may slow growth or progression. Recently, the gut microbiome has been implicated in a wide-range of roles in tumour biology. Through modulation of immunity, the gut microbiota can improve the efficacy of several immunotherapies. However, despite the prevalence of breast cancer, there is still a lack of microbiota studies in this field, including exploring the influence of external microbiome-modulating factors such as antibiotics. We describe herein how disruption of the gut microbiota via antibiotics may be detrimental to patient outcomes through acceleration of tumour growth.

**Results:** Supplementing animals with a cocktail of antibiotics leads to gut microbiota alterations and is accompanied by significant acceleration of tumour growth. Surprisingly, and distinct from previous microbiome-tumour studies, the mechanism driving these effects do not appear to be due to gross immunological changes. Analysis of intratumoural immune cell populations and cytokine production are not affected by antibiotic administration. Through global tumour transcriptomics, we have uncovered dysregulated gene expression networks relating to protein and lipid metabolism that are correlated with accelerated tumour growth. Fecal metabolomics revealed a reduction of the microbial-derived short-chain fatty acid butyrate that may contribute to accelerated tumour growth. Finally, through use of a routinely administered antibiotic in breast cancer patients, Cephalexin, we have shown that tumour growth is also significantly affected. Metataxanomic sequencing and analysis highlighted significant antibiotic-associated reductions in the butyrate producing genera *Odoribacter* and *Anaeotruncus*, and increased abundance of *Bacteroides*.

**Conclusions:** Our data indicate that perturbation of the microbiota by antibiotics may have negative impacts on breast cancer patient outcomes. This is of importance as antibiotics are regularly prescribed to breast cancer patients undergoing mastectomy or breast reconstruction. We have also shown that the metabolic impact of disruption to the microbiome should be considered alongside the potent immunological effects. We believe our work lays the foundation for improving the use of antibiotics in patients, and with further investigation could potentially inform clinical practice.

## Background

Breast cancer (BrCa) is the second most prevalent cancer globally and the most prevalent in women [1]. It was estimated to contribute 11.6% of the 18.1 million new cancer diagnoses and 6.6% of the 9.6 million cancer related fatalities in 2018 [1]. While ~10% of BrCa cases are linked to hereditary or somatic mutations in tumour suppression genes, such as BRCA1 and BRCA2, the vast majority of cases are the result of lifestyle and environmental factors [2]. Hormone therapies, smoking, alcohol consumption, and diet have all been associated with the onset of BrCa, the latter likely being linked to a disruption in gut homeostasis [1–3].

The gut microbiota comprises a diverse and complex array of microbes which play an integral role in maintaining human health. Under normal healthy conditions, these microbes regulate our immune system through molecular interactions at the intestinal barrier [4]. However, when the gut environment is altered unfavourably, such as after a course of antibiotics, the microbial community profile is shifted or disturbed, and gut homeostasis is lost [5, 6].

Alterations in the gut microbiota are associated with an array of molecular and physiological changes. Inflammatory signalling pathways can be amplified or dampened depending on changes in bacterial metabolite production, and such alterations have been associated with a variety of diseases, including cancer [7, 8]. In colorectal cancer, a reduction in short-chain fatty acid (SCFA) production by *Roseburia* resulted in a proinflammatory cascade that promoted cancer progression in an *in vivo* mouse model [9]. Contrastingly, inoculation of mice harbouring subcutaneous melanomas with *Bifidobacterium*, a known beneficial or ‘probiotic’ genus, has been shown to amplify the anti-tumour effect of an anti-PD-L1 immunotherapy through the priming of CD8^+^ T-lymphocytes [10]. Studies like these demonstrate the microbiota’s integral role in regulating local and systemic responses to cancer.

Since the discovery of penicillin in 1928, antibiotics have become an extremely effective way of preventing and fighting bacterial infections [11]. Nevertheless, with the evolution of antibiotic-resistant bacterial strains, and an emerging understanding of the risks associated with antibiotic-induced microbiota disturbances, the ubiquity of their use has become increasingly controversial [11, 12]. Whilst the use of prophylactic antibiotics to treat BrCa patients, particularly following a mastectomy and reconstructive surgery, is common practice, their clinical benefit is under debate [13, 14]. Additionally, the resultant alterations in the gut microbiota created by their use raises concerns over potential impacts on metabolism and inflammation that might drive tumorigenesis [15, 16]. Whilst the consequence of antibiotic use has been studied in other cancers, the impact on BrCa is largely unknown.

This research aimed to identify how antibiotic-induced gut microbiota changes influences the progression of BrCa. Using MMTV-PyMT derived PYMT-BO1 and spontaneous EO771 breast carcinoma cells in an orthotopic, mammary fat pad injection model, we identified a significantly increased rate of primary tumour growth in animals subjected to a broad-spectrum cocktail of antibiotics. Importantly, this also occurred in Cephalexin-treated animals, an antibiotic commonly administered to BrCa patients post-surgery in the US [16]. Immunological and proteomic analysis found little variation in the immune cell infiltration of tumours and no change in tumour cytokine production in antibiotic treated animals. However, whole tumour transcriptomics identified profound differences in regulation of metabolism pathways. Subsequent metabolomic analysis of fecal material following antibiotic treatment highlighted significant changes in SCFA metabolites, particularly the reduction of butyrate and acetate in addition to changes in microbiota derived amino acid production. Additionally, analysis of the microbiome in Cephalexin treated animals showed a significant reduction in butyrate-producing bacteria.

These findings highlight a possible mechanism by which antibiotic administration may be detrimental to patient outcomes. While it is not feasible to rule out their use, further consideration must be taken to optimise their use in a clinical setting. It is hoped that with further studies our work may help guide clinical practice through use of better targeted antibiotics in BrCa patients.

## Results

### Treatment with broad spectrum antibiotics results in severe perturbation of the gut microbiota and acceleration of breast tumour growth

To investigate the role of the microbiota on breast tumour growth we employed an orthotopic mammary fat pad injection model using two distinct cell lines. The PyMT-BO1 line is derived from MMTV-PyMT animals and exhibits a luminal B intrinsic phenotype, whilst the EO771 line is derived from a spontaneous C57BL/6 breast tumour, and more closely resembles basal BrCa [17]. Prior to tumour cell injection, the microbiota of animals was depleted using a cocktail of antibiotics consisting of Vancomycin, Neomycin, Metronidazole and Amphoteracin (VNMA) via oral gavage with Ampicillin also available in drinking water. This cocktail has been previously shown to result in severe microbial changes [18, 19]. Following the regimen documented in Figure 1A, animals treated with the VNMA cocktail had very low fecal DNA concentrations, compared to water treated controls (Data not shown). Samples which could be quantified were subjected to PCR amplification of the V1+2 region of the 16S rRNA gene. Amplifiable DNA is present in all starting samples in both groups, however, after 5 days of VNMA treatment, amplifiable DNA is no longer present in feces and is maintained for at least 22 days of treatment (DoT) (Figure 1B) indicating a severe microbiota knock-down in these animals. Crucially, animals with a depleted microbiota were shown to have significantly accelerated tumour growth when compared to water treated counterparts, both in the PyMT-BO1 (Figure 1C and D) and the EO771 model of BrCa (Figure 1E). To determine any phenotypic differences in tumour properties, we used H&E staining to examine tumour architecture, however no significant changes were observed in antibiotic treated vs. control animals (Figure 1F).

**Figure 1.**
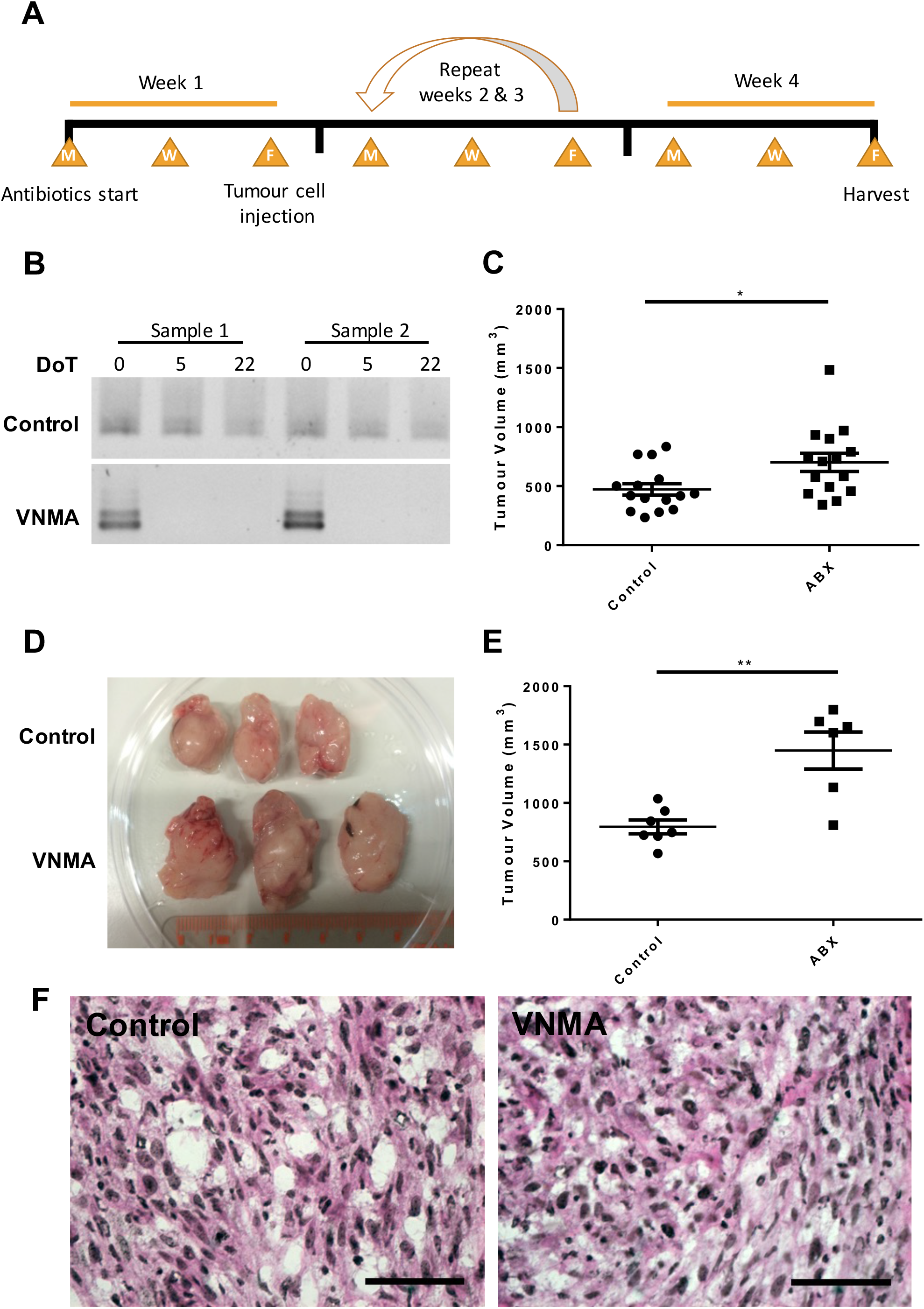
Tumour growth is accelerated by VNMA administration: A) Schematic showing antibiotic treatment regimen, (M – Monday, W – Wednesday and F – Friday). B) Representative images of agarose gels used to detect 16S rRNA amplification by PCR of DNA extracted from fecal samples. DoT – Days of Treatment. C) *Ex vivo* tumour volumes from animals injected with the PyMT derived PYMT-BO1 tumours. D) Representative photograph showing size of control and VNMA treated tumours. E) *Ex vivo* tumour volumes from animals injected with the spontaneously derived EO771 cell line. F) Representative images of H&E staining of tumours from control (Left) and antibiotic (Right) treated animals. Scale bar = 50μm, significance determined by unpaired, two-sample t test, * p<0.05 ** p<0.01, n=15 for PYMT-BO1 and n≥6 in EO771.

### Antibiotic-induced microbiota changes do not alter frequencies of infiltrating leukocytes, myeloid cells, or to macrophage polarisation

Given the known interactions between the microbiota and the immune system, and the previously published literature suggesting that manipulating the microbiota can alter antitumour immunity, we decided to undertake high level immune cell phenotyping using flow cytometry. We performed profiling of intratumoural CD11b^+^ myeloid cells, and subsequently analysed proportions of F4/80^+^ macrophages and Ly6G^+^ neutrophils. We did not observe any significant changes in either population (Figure 2A). In addition, we profiled the activation state of tumour-associated macrophages (TAMs) using MHC-II and CD206 to delineate “M1” and “M2” polarisation. However, we did not observe any significant differences in either the number of cells presenting these markers or the median fluorescence intensity (MFI) (Figure 2A&B). Whilst the proportion of immune cells in the tumour was overwhelmingly weighted towards myeloid cells (90-95% of all CD45+ events), we also undertook profiling of T cell populations, identified as CD3+CD4+ and CD3+CD8+. T regulatory (T_reg_) cells were also identified from CD4+ cells by FoxP3 staining (Figure 2C). This analysis was also performed in spleen and mesenteric LN as a measure of peripheral immune cell populations, however no changes were observed at the tumour or in either organ (Figure 2D&E). After determining that immune cell populations did not appear to be significantly altered after VNMA treatment, we undertook analysis of intratumoural cytokine production by multi-plex assays. However, using whole tumour protein lysates we were unable to detect any significant changes to intratumoural cytokine production (Figure 2F). Conversely, cytokine analysis of colon tissue revealed multiple cytokines were significantly reduced by VNMA, including CXCL-1 IL-1β, IL-2 and TNF-α (Supplementary Figure 1).

**Figure 2.**
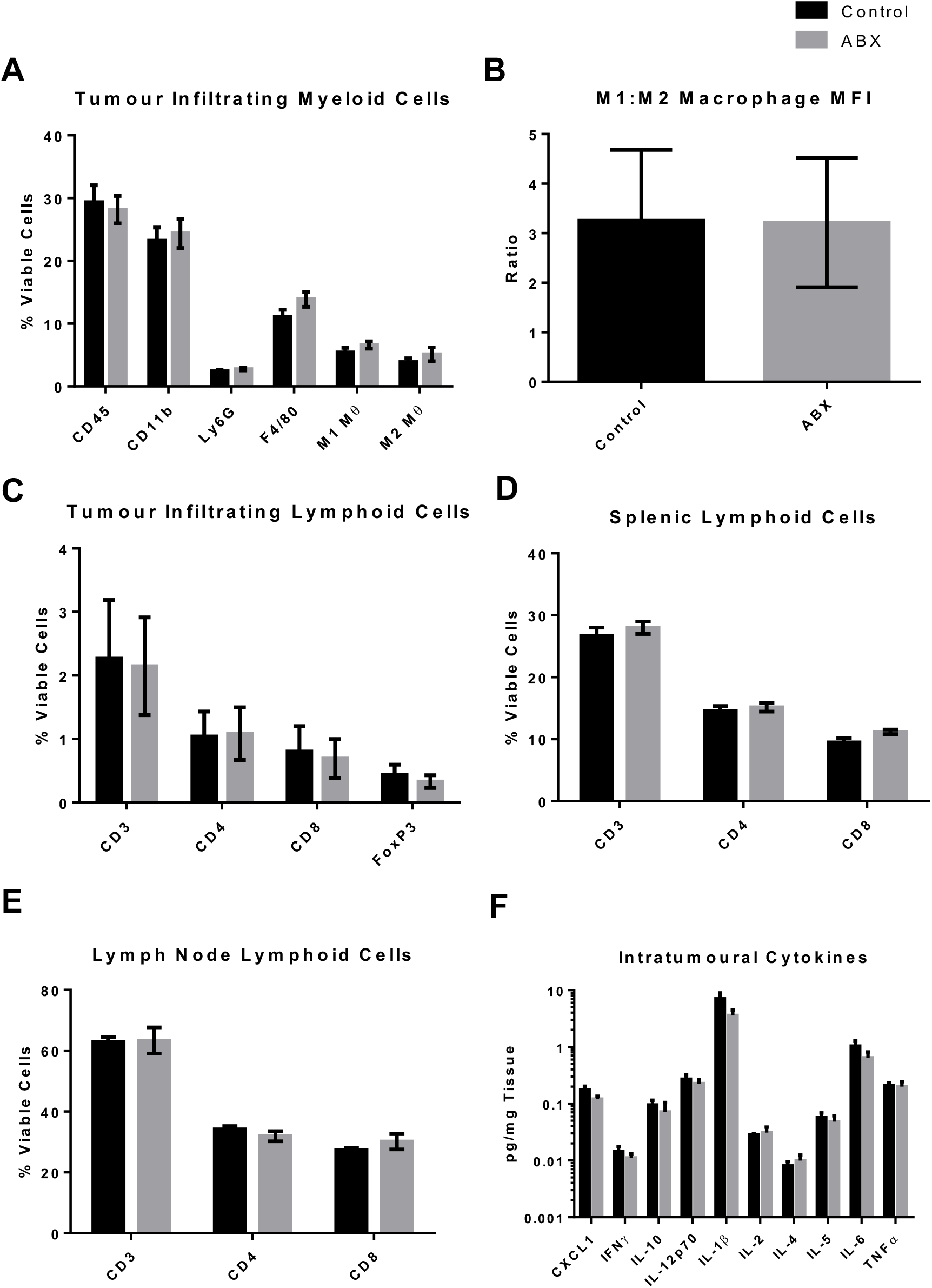
Immune cell populations are not altered by VNMA administration: A) Myeloid cell populations determined by flow cytometry described as a percentage of total live cells in the tumour. B) Median Fluorescence Intensity analysis of CD206 and MHCII in macrophages. C) Flow cytometry of tumour infiltrating T cells, CD45 was again used as a pan-leukocyte marker (not shown). D&E) Flow cytometric analysis of spleen and mesenteric lymph nodes. F) Intratumoural cytokine production analysed by MSD V-Plex assays, data is shown in logarithmic scale to account for large intra-cytokine differences in production.

### Transcriptomic analysis of whole tumour RNA reveals a gene expression pattern consistent with changes to metabolic processes

The lack of changes at an immunological level led us to question what might be driving our phenotype. To address this, we undertook global transcriptomic sequencing of whole tumour RNA using 3 samples from each experimental condition. This differential gene expression analysis yielded a total of 172 differentially expressed genes (DEGs): 85 upregulated and 87 downregulated in the VNMA treated animals, with respect to controls. The top 7 of each are annotated in the volcano plot (Figure 3A). The entire DEG list is available in Supplementary Figure 2. These genes were used for functional clustering analysis in the NIH DAVID tool according to Gene Ontology (GO) biological process definitions. The full list of biological processes highlighted using this method is available in Supplementary Tables 1 and 2, however a high frequency of processes involved in cellular metabolism were identified. In total, 91 of 172 DEGs were associated with metabolic transduction, transcription, and migration and differentiation (Figure 3B). Given the overwhelming prominence of metabolic gene regulation in our analysis, we chose to delve further into the biological functions that may be consequentially altered. Using lower level GO definitions, we observed significant changes in lipid metabolism, gluconeogenesis, and protein metabolism (Figure 3C). Further analysis of these biological functions revealed two major groups of genes. One belonging to lipid metabolism, such as *ACACB, LPL* and *ACSL1*, whilst the other contains several genes relating to protein modification or metabolism such as *FBXL5, MADD, OAZ2* and *TIPARP* (Figure 3C&D). Genes associated with apoptosis, cell signalling, migration and differentiation are also significantly enriched in our DEGs and are described in Supplementary Figure 3.

**Figure 3.**
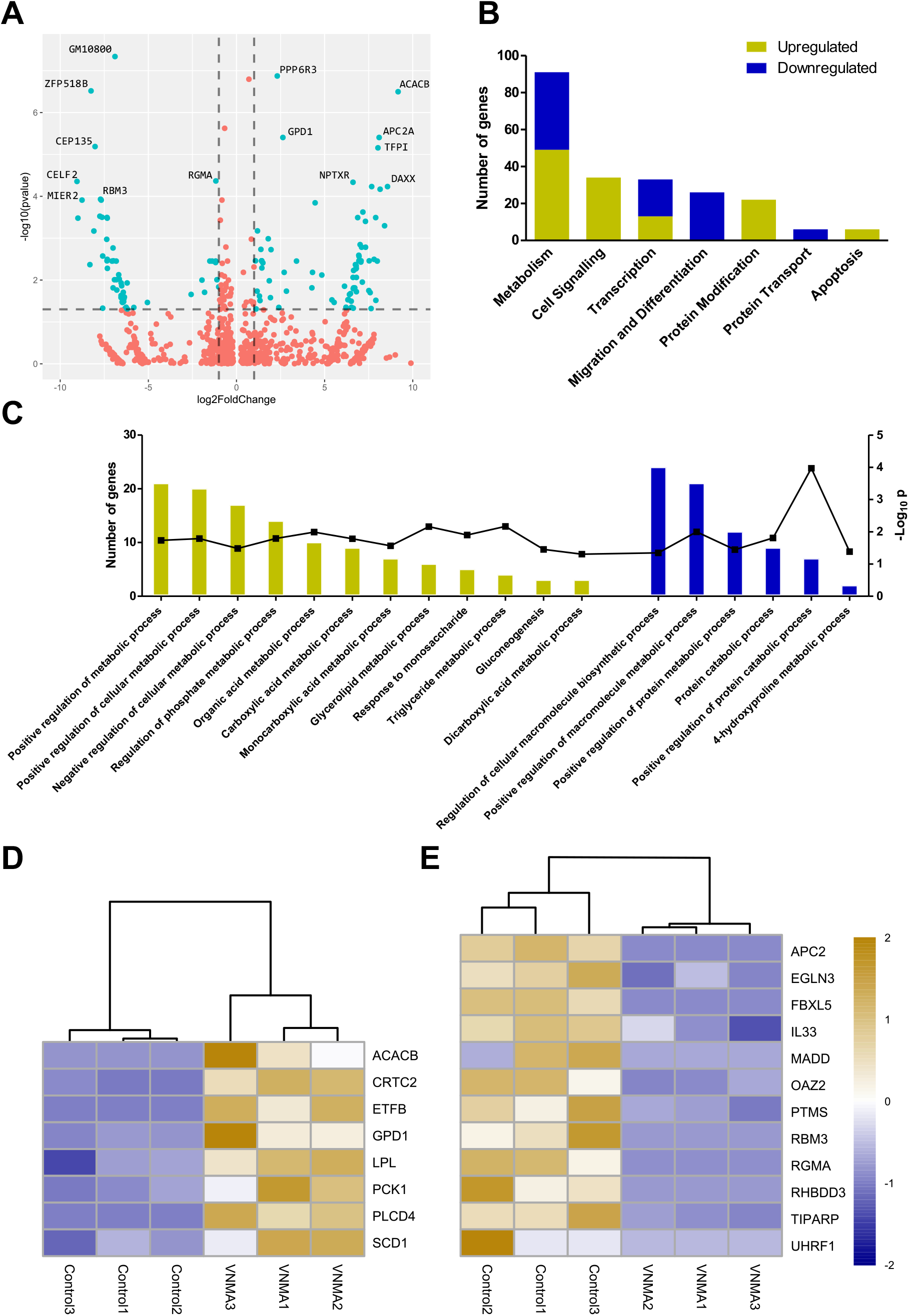
Intratumoural gene regulation is significantly different after VNMA treatment, particularly in metabolic processes: A) Volcano plot describing the parameters used for differential expression, FDR adjusted p value < 0.05 (Log_10_ adjusted) and Fold Change >2 (Log_2_ adjusted). The top 7 DEGs are annotated on the graph. B) High level analysis of biological process enrichment using DAVID, separated by over-arching biological function C) Low-level analysis of the metabolic processes enriched in our DEG set, data is shown according to the number of genes involved in each process (Bar-plot, Left Y-axis) and the significance of the enrichment (Line, Right Y-axis, Log_10_ adjusted). D&E) Heatmaps showing specific genes which are related to lipid (D) and protein (E) metabolism from our DEG set. Colour ratio is shown according to Log_2_ fold change.

### Metabolomic analysis of antibiotic-treated fecal samples indicates profound alterations in microbially-derived metabolites

As our transcriptomic analysis suggested antibiotic-induced acceleration of BrCa tumour growth may be related to metabolic reprogramming, we further probed this relationship via metabolomic analyses. Given we are unable to obtain information regarding the constituents of the microbiota in antibiotic treated animals (due to very low DNA yield), we decided to perform fecal metabolomics as a proxy. Fecal water samples were used in ^1^H NMR analysis which revealed that metabolite production in the antibiotic treated animals was significantly altered when compared to control animals as indicated in PCA (Figure 4A). Further analysis of metabolites revealed 17 which were significantly different, 8 were elevated whilst 9 were significantly depleted after antibiotic treatment (Figure 4B). The individual metabolites responsible for these changes are described in Figure 4C. Several amino acids including alanine, histidine and aspartate are significantly increased in the antibiotic treated animals in addition to the fermentation substrate and product raffinose and lactate respectively. Conversely, the short chain fatty acids butyrate and acetate are significantly decreased by antibiotic administration in addition to the branched chain fatty acid isovalerate (Figure 4C).

**Figure 4.**
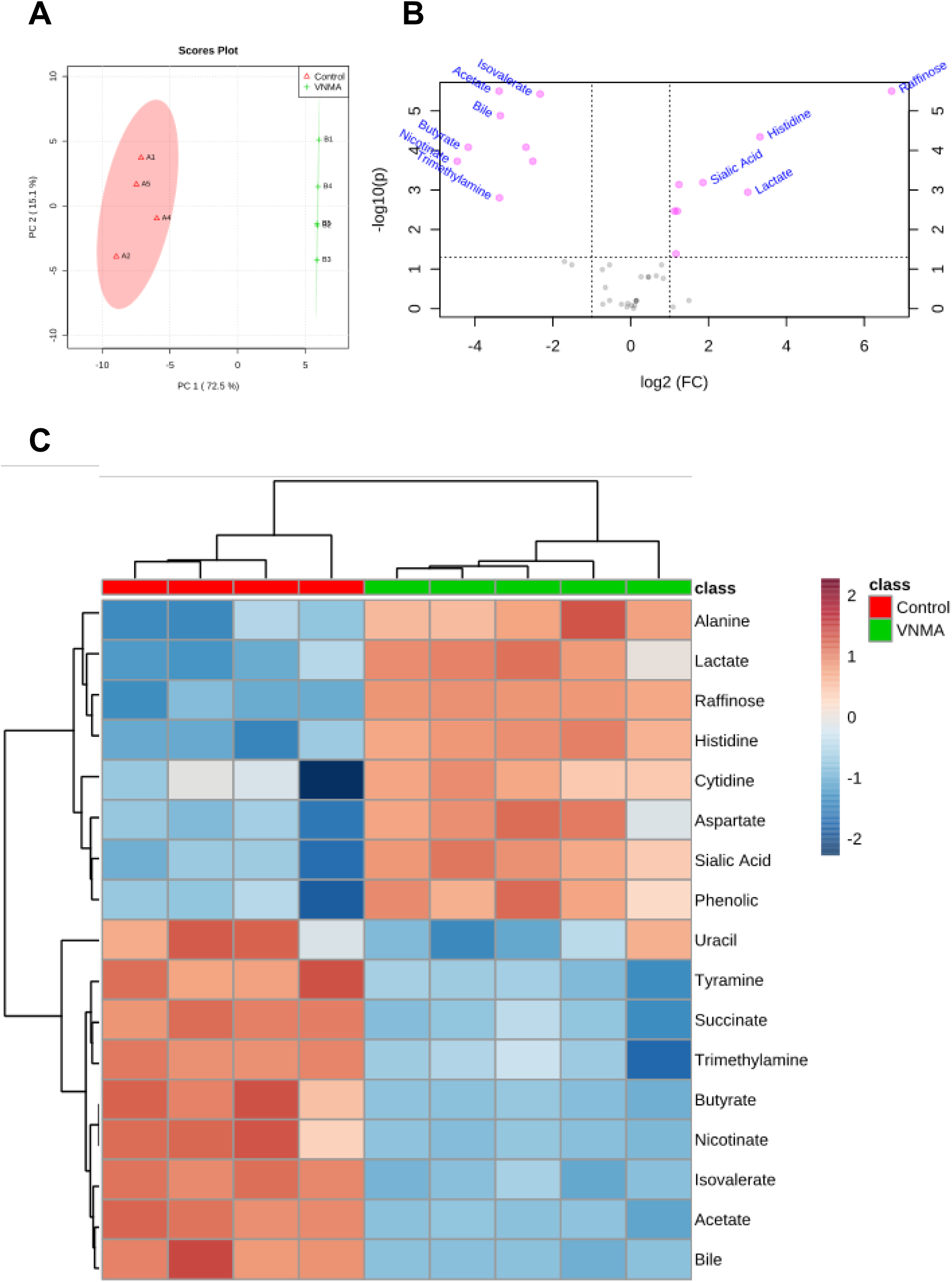
Antibiotic administration results in changes to bacterial metabolite production: One biological replicate had significant outliers in multiple metabolites and was excluded from analysis but is documented along with the full metabolomic dataset in Supplementary Figure 5. A) Two component PCA of included biological replicates. B) Volcano plot of remaining replicates; x-axis specifies the Log2 fold change of VNMA relative to control animals and the y-axis specifies the negative logarithm to the base 10 of the t-test p-values. Dashed lines represent cut off values for differential regulation (FC ± 1, padj < 0.05). The 10 most significantly regulated metabolites are annotated. C) Filtered heatmap clustered by average Euclidean distance showing only significantly regulated analytes (p<0.05), calculated by unpaired, two-tailed t test. All graphs produced using MetaboAnalyst 3.0 software.

### Treatment with a single, BrCa relevant antibiotic also results in acceleration of tumour growth and causes significant changes to the microbiota

Due to the knock-down efficacy of the previously used VNMA antibiotic cocktail, we were unable to amplify microbial 16S rRNA DNA for sequencing and were therefore unable to profile the microbiota of these animals. In addition, whilst clinically relevant in certain scenarios, the VNMA cocktail is not applicable to most BrCa patients. We therefore turned our attention to BrCa relevant antibiotics, particularly Cephalexin, which is widely prescribed to BrCa patients in the US after mastectomy. Treatment of C57BL/6 animals harbouring PYMT-BO1 tumours with Cephalexin at a patient relevant dose (8.64mg/kg) led to significantly accelerated tumour growth (Figure 5A). Analysis of the microbiota of these animals revealed that, post-treatment, the antibiotic and control animals cluster independently of their pre-treatment samples. Additionally, the post-treatment samples of the control and antibiotic treated samples also cluster differently (Figure 5B). The changes in abundance that lead to this clustering appear to be related to an interplay between the genera *Lactobacillus* and *Faecalibaculum*. Visualisation of relative abundance by bar plot shows that both the control and antibiotic treated animals lose relative abundance of Lactobacilli over time. In the control animals, this ‘gap’ appears to be predominantly replaced by *Faecalibaculum*, however this does not occur in the antibiotic treated animals. Instead, several other genera have increased relative abundance (Figure 5C). This change is highlighted by alpha diversity analysis that shows the reciprocal Simpson index significantly increases from 2.352 in the pre-treatment samples to 5.422 after Cephalexin treatment. By comparison, whilst the microbiota of control animals also increases in diversity during experimentation, this is non-significant (2.486 to 3.073) (Figure 5D). Further analysis revealed that 11 genera were differentially abundant. When accounting for multiple comparisons, 8 genera were significantly altered by antibiotic treatment: *Mucisprillum, Marvinbryantia, Parabacteroides, Anaeroplasma, Bacteroides* and *Paraprevotella* were significantly increased, whilst *Alloprevotella, Alistipes, Odoribacter, Faecalibaculum and Anaerotruncus* were significantly decreased after Cephalexin treatment (Figure 5E).

**Figure 5.**
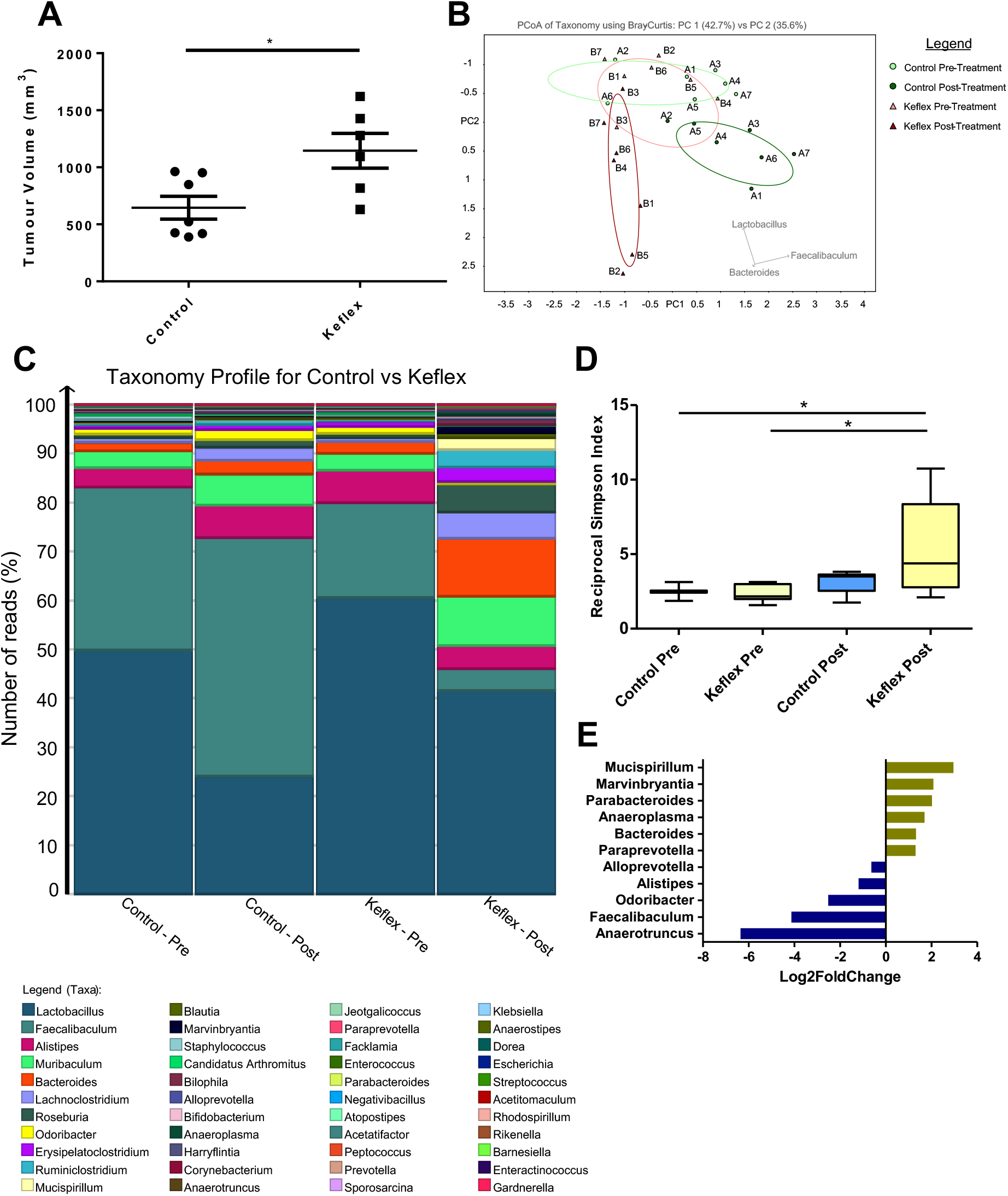
Administration of Cephalexin increases tumour growth and 16S rRNA analysis describes the related changes: A) Tumour volumes of animals implanted with PYMT-BO1 tumours treated with patient analogous doses of the BrCa relevant antibiotic Cephalexin. Significance determined by unpaired two-sample T-test, * p<0.05, n≥6 B) Principle component analysis of all replicates using Bray Curtis distances. Pastel colours represent starting microbiome and darker are end-point samples. C) Full microbiome composition by genera in control vs Cephalexin treated animals at experimental start and endpoints. Bars represent percentage of total reads for each genus and are sorted by increasing number from bottom to top. Legend is ordered by greatest abundance. D) Alpha diversity analysis by reciprocal Simpson index of control vs Cephalexin treated animals at pre- and post-treatment timepoints, significance was determined by one-way ANOVA using Tukey’s test for multiple comparisons, * p<0.05, n=7. E) Mean fold change of significantly altered genera in the antibiotic treated animals relative to control at the post-treatment sampling point. Yellow indicates genera which are significantly enriched whilst blue bars are depleted in Cephalexin treated animals. Significantly altered genera were determined by unpaired two-sample or paired T test with FDR correction of 5%.

## Discussion

The use of antibiotics is widespread amongst cancer patients to prevent opportunistic infection during periods of immunocompromisation. However, now more than ever is the time to reevaluate antibiotic use in the clinic. The threat of antibiotic resistant pathogens is imminent and serious. According to the 2016 review by the UK Department of Health, antimicrobial resistance is already killing 700,000 people per year worldwide and this figure is expected to grow exponentially over the next 30 years [20]. Additionally, some evidence suggests that antibiotic use may not be beneficial to all patients. Recent studies have demonstrated an unequivocal role of the patient microbiome in orchestrating anti-tumour responses, and many have found that the use of antibiotics compromises treatment efficacy in several cancers. It is therefore prudent that clinicians begin to carefully consider the efficacy of antibiotic use in their patients. To do so, we must fully understand how the microbiome impacts different cancer pathologies. Other groups have made progress in understanding how antibiotics affect immunogenic cancers, however there is a paucity of data regarding how the microbiota might impact BrCa [21–23].

Our work intended to understand whether the use of antibiotics has any impact on primary tumour growth in BrCa. To address this, we undertook tumour studies using orthotopically implanted PyMT derived luminal (PYMT-BO1 or spontaneously derived basal (EO771 tumours in animals which had been administered a robust, VNMA antibiotic cocktail. This revealed that severe disruption of the gut microbiota results in accelerated tumour growth across both models. Importantly, this suggests that antibiotic treatment is detrimental regardless of the BrCa intrinsic subtype, however we are yet to test this in additional subtypes. This is largely in agreement with the findings of groups studying other cancers. Use of antibiotics has been shown to impact tumour growth in both pre-clinical, and human studies. However, these studies focus on the influence of the microbiome on anti-tumour therapies. For example, Vetizou *et al*. and Routy *et al*. probe the impact of antibiotics on anti-CTLA4 and anti-PD-1 therapies respectively, finding that these treatments are rendered ineffective when the microbiome is depleted [23, 24]. However, when comparing control and antibiotic-treated animals without administration of anti-tumour agents, these groups found no difference in tumour volume. This suggests that our findings may be specific to BrCa, and is supported by Rossini *et al*., who in 2006 demonstrated, using HER2/*neu* transgenic mice, that antibiotic administration alone increased the incidence of spontaneous BrCa [25]. Analysis of our tumours by H&E staining revealed no overt differences in tumour architecture or cellular makeup between control and antibiotic tumours. Therefore, based on the known roles of the gut microbiota in guiding anti-cancer immune responses, we hypothesised that the mechanism driving our phenotype was likely to be immunological. However, after wide profiling of immune cell populations at the tumour and peripherally, we were unable to find any significant differences. We also undertook analysis of cytokine production using MSD V-PLEX assays to quantitatively assess intratumoural and intestinal cytokine production. Whilst none of the profiled cytokines were significantly altered intratumourally, several were significantly decreased in intestinal tissue. Both IL-1β and TNFα play key roles in orchestration of gut-immune responses through chemoattraction and modulation of inflammation. Both have also been shown to drive CXCL1 expression and their decreased production may explain our observed significant reduction of CXCL1 in intestinal tissues following VNMA treatment [26, 27]. The biological impact of this cytokine dysregulation on tumour growth is unclear, however it is suggestive of disrupted gut homeostasis in the VNMA treated animals.

In the absence of immunological changes, we employed RNA sequencing of whole tumour extracts to gain further mechanistic insight. In agreement with the lack of apparent immune infiltration into tumours, we determined, after biological process enrichment analysis, that alterations were predominantly seen in metabolic processes, particularly in lipid and protein metabolism. Metabolic reprogramming is a well-established hallmark of cancer and upregulation of lipid metabolism is strongly associated with tumorigenesis, particularly in BrCa. The significant upregulation of lipoprotein lipase (LPL) expression in our model suggests increased utilisation of circulating fatty acids. In support of this, Acyl-CoA Synthetase 1 (*ACSL1*) is also upregulated in tumours. Its role is to activate long chain fatty acids during synthesis of acyl-CoA and is the first committed step in fatty acid metabolism. Upregulation of both *ACSL1* and *LPL* expression has been demonstrated in BrCa and correlate negatively with overall survival [28–30]. In addition to meeting the cell’s energy demands, lipid metabolism also provides substrates for membrane lipid generation. A key enzyme in this process, Stearoyl-CoA desaturase (SCD1) is also significantly upregulated after VNMA administration. Its primary function is catalysing the production of monounsaturated fatty acids (MUFAs) from their saturated counterparts. Production of MUFAs is noted to be elevated in several cancers, including breast, and is required for generation of cellular lipids, particularly membrane phospholipids [31]. Additionally, elevated SCD1 levels prevent the accumulation of saturated fatty acids, which can induce apoptosis. This is demonstrated by SCD1 inhibition in BrCa cells which demonstrate decreased proliferative ability and increased rates of cell death. Furthermore, high levels of SCD1 in BrCa are associated with poor prognosis [32, 33]. In addition to changes in lipid metabolism, several genes associated with protein metabolism are also downregulated in our DEG set, and many of these genes are known tumour suppressors. Expression of BNIP3 can induce apoptosis in response to oxidative stress by mitophagy and is seen at reduced levels in more aggressive BrCa subtypes [34, 35]. Additionally, Ornithine Decarboxylase Antizyme 2 (OAZ2) has been shown to inhibit the enzyme ornithine decarboxylase (ODC) which synthesises polyamines from ornithine and promotes proliferation in numerous cancers [36, 37]. The network of gene expression identified by our transcriptomic analysis represents several potential avenues by which antibiotic administration may be accelerating BrCa growth. However, robust metabolomic analyses are required to confirm these observations and should be the focus of future investigation.

The question remains how could perturbation of the microbiome induce these tumour-enhancing metabolic effects? Microbiota-derived metabolites have previously been shown to alter tumour growth in other models and recently it has been demonstrated that this is also applicable to BrCa. Administration of cadaverine to mice harbouring 4T1 BrCa tumours resulted in improved outcomes (smaller tumours, fewer metastases). Furthermore, cadaverine production was shown to be decreased in BrCa patients and correlated with survival [38]. To test this in our BrCa models we used fecal metabolomic analysis by NMR and observed significant metabolic dysregulation in the microbiota of VNMA treated animals (Figure 4). Of note are perturbations in the SCFAs acetate and butyrate. Gut microbiota derived butyrate is readily absorbed and has been suggested to play a role in multiple diseases including cancer through its ability to inhibit histone deacetylases (HDACs). Inhibition of HDACs by butyrate has been shown to sensitise cancer cells to ROS induced apoptosis [39]. *In vitro* administration of exogenous butyrate suppresses proliferation of BrCa cells through senescence and induces apoptosis [40]. Whilst our transcriptomic data did not reveal any differences in transcription of cell cycle regulatory genes, we do observe decreased expression of pro-apoptotic genes such as *BNIP3* and increased pro-survival genes such as *HERPUD1* and *URI* which is consistent with butyrate’s bioactivity. Therefore, we hypothesize that decreased butyrate bioavailability is playing some role in our system.

Several compounds that can be utilised by tumours were also upregulated in VNMA treated animals. Both alanine and lactate have been associated with metabolic reprogramming, however alanine alone is not sufficient to contribute to the increased energy demands of tumours [41]. However lactate achieves this by feeding into the Krebs cycle via conversion into pyruvate [42]. Additionally, lactate has HDAC inhibitor activity and may modulate gene expression in a similar fashion to that of butyrate [43]. Though the implications of this on tumorigenesis are currently unclear, the effect of microbiota derived lactate in BrCa is under investigation.

To strengthen the clinical relevance of our data and attempt to gain some insight into microbial population changes, we turned to using a BrCa relevant antibiotic regimen. Cephalexin is frequently prescribed to BrCa patients in the US undergoing mastectomy and treating animals at a patient relevant dose (8.64mg/kg) resulted in significantly accelerated tumour growth. Profiling of the microbiota revealed significant changes to the microbial constituents of Cephalexin treated animals. The most significantly changed genus in both treatment groups relative to the pre-treatment sample was *Lactobacillus*, however there was no significant difference between treatments. Therefore, the loss of *Lactobacillus* is likely driven by cage effects, tumour-microbiota interactions or natural maturation of the microbiome, and is not a result of Cephalexin administration [44, 45]. Of the genera significantly depleted relative to the control, several are either known butyrate producers (i.e. *Odoribacter* and *Anaerotruncus*) or possess the genes necessary for butyrate production *(Faecalibaculum* and *Alistipes*) [46–48]. This is consistent with the significantly decreased butyrate production observed in our fecal metabolomics. Whilst microbiota derived butyrate has been shown to inhibit colorectal cancer cell proliferation *in vitro*, tumour studies have demonstrated a paradoxical effect [49]. As discussed previously, butyrate has been shown to induce apoptosis in BrCa cultures, but the effect *in vivo* is yet to be characterised and is under investigation.

Of the increased relative abundance genera, most are understudied with respect to their contribution to human health. Of particular interest is the significant increase in *Bacteroides* spp. which are known to be resistant to several antibiotics including β-lactams [50]. Their role in human health and particularly contributions to tumorigenesis is mixed, and heavily species/strain dependent. An abundance of *Bacteroides fragilis (B. fragilis)* has been shown to potentiate immunotherapy in mouse sarcoma [23]. However, their presence in the microbiome has recently been associated with increased incidence of colon carcinogenesis [51]. Additionally, different species have also been shown to drive distant cancers. *Bacteroides thetaiotaomicron* is associated with non-response to PD-1 therapies in melanoma through reduced infiltration of cytotoxic T cells [21]. Therefore, to determine the potential importance of *Bacteroides* in our system, it is essential we obtain data from metagenomic analyses to determine the species changes in the microbiome of Cephalexin treated animals; these studies are currently underway.

Our work has shown that disruption of the gut microbiota using robust antibiotics may have detrimental impacts on BrCa growth. Whilst the mechanism driving these effects is currently unclear, we have uncovered a potential network of metabolic processes that may be contributors. Furthermore, we have shown that antibiotic administration leads to dysregulation of bacterial metabolite production that has the potential to release the ‘brakes’ on tumour growth. Finally, using a BrCa relevant antibiotic we have shown that BrCa growth is also accelerated, suggesting even small perturbations of the microbiome may impact patient outcomes. We have demonstrated that this is associated with a loss of butyrate producing genera in the microbiome and increases in several other genera. Whilst it is too early to say the use of antibiotics in the clinic should be reconsidered, we believe it is necessary to investigate these findings further to ensure patient outcomes are not impacted by antibiotic use.

## Materials and Methods

### Animals

All animals were female C57BL6 mice and were sourced in-house. All animals were age matched at 8-10 weeks old and were cage mixed prior to experiments. All animal experiments were performed in accordance with UK Home Office regulations and the European Legal Framework for the Protection of Animals used for Scientific Purposes (European Directive 86/609/EEC).

### Antibiotic Administration

Animals were treated with antibiotics 3 times weekly by oral gavage (200μl in water). Animals were treated with either an antibiotic cocktail consisting of 1mg/ml Amphoteracin B (Sigma-Aldrich, St-Louis, Missouri, USA), 25mg/ml Vancomycin (Sigma), 50mg/ml Neomycin (Sigma), 50mg/ml Metronidazole (Sigma) with drinking water being supplemented with 1mg/ml Ampicillin (Sigma) or 44mg/ml Cephalexin (Sigma). Antibiotic treatment began 5 days prior to tumour cell injection and was maintained throughout animal experiments.

### Breast cancer cell culture

PYMT-BO1 and EO771 cells were cultured in high glucose DMEM (Invitrogen, Carlsbad, California, US) supplemented with 10% fetal bovine serum (FBS) (Hyclone, Invitrogen) and 100 units/mL penicillin/streptomycin (Pen/Strep) (Invitrogen). Cells were maintained at 37°C and 5% CO_2_. Tissue culture plastic was coated with 0.1% porcine gelatin (Sigma) in water for 1 hour at 37°C prior to culture.

### *In vivo* tumour growth assays

Syngeneic mouse breast carcinoma (PYMT-BO1^a^ or E0771^b^) cells were injected at 1×10^5^ per 50μl of a 1:1 mixture of PBS and Matrigel (Corning Life Sciences, Corning, New York, USA) into the left inguinal mammary fat pad (MFP) of age matched female mice. Tumours were measured in two dimensions (Length x Width) every two days from 7 days post injection (DPI) using digital calipers. Upon conclusion of the experiment or once the tumours reached 1000mm^3^ the animals were humanely killed by cervical dislocation and tissues harvested for various downstream analyses. Tumour volume was calculated according to the following formula: length * width^2^ * 0.52.

### Paraffin embedding and sectioning of formalin fixed tissues

Tissues were fixed overnight in 4% PFA at 4°C before washing twice in PBS for 30 minutes. Tissues were gradually dehydrated through successively increasing concentrations of ethanol (30%-100%) before clearing in Histoclear (National Diagnostics, Atlanta GA, USA) and embedding in paraffin. Paraffin blocks were sectioned using a HM355 S microtome (Microm, Bicester, UK) at a thickness of between 5μm and 10μm and mounted onto positively charged glass slides (Thermofisher). Sections were dried o/n at 37°C. Prior to staining sections were rehydrated by washing in Histoclear before incubations in gradually decreasing concentrations of ethanol (100%-50%) with a final wash in dH_2_O.

### H&E staining

Frozen tumour sections were air dried for 10 minutes at room temperature, transferred to running tap water for 30 seconds then placed in Mayer’s Hematoxylin for five minutes. Sections were rinsed in running tap water until blue, drained and Eosin was added for 20 seconds. Excess Eosin was blotted off sections, and sections were given a quick rinse in running tap water followed by a graded dehydration to Histoclear. Sections were mounted with DPX and allowed to air dry.

### Flow Cytometry

Organs were excised from humanely killed animals and tissues were mechanically homogenised using scalpels. Homogenate was incubated in collagenase solution (0.2% Collagenase IV (Invitrogen), 0.01% Hyaluronidase (Sigma) & 2.5U/ml DNAse I (Sigma) in HBSS) for 1 hour at 37°C with regular agitation. Supernatant was passed through a 70μm cell strainer and centrifuged for 5 minutes at 300 x g/4°C. Pellet was washed twice in PBS and resuspended in 10ml 1X red blood cell lysis buffer (Invitrogen) and incubated for 5 minutes at RT. Cells were washed once in PBS, counted using a haemocytometer (Sigma) and 1 million cells per condition transferred to a 96 well plate for staining. Cells were incubated in a fixable Live/Dead stain (Invitrogen, Thermofisher) for 30 minutes at RT, washed twice and blocked in Fc Block (Miltenyi, Bergisch Gladbach, Germany) made in FACS buffer (1% FBS in PBS) for 10 minutes at 4°C. Cells were resuspended in 100μl antibody solutions (Table 1) and incubated at 4°C for 30 minutes in the dark. For cell surface only staining, cells were incubated in 4% PFA for 30 minutes, washed once in PBS and stored at 4°C until analysed. If intracellular staining is required, cells were incubated in FoxP3 fixation/permeabilisation buffer (Thermofisher) overnight at 4°C, washed twice in 1X permeabilisation buffer (Thermofisher), blocked in 5% normal rat serum for 30 minutes at RT and stained in the relevant antibody diluted in 1X permeabilisation buffer for 30 minutes at RT in the dark. Cells were washed twice in 1X permeabilisation buffer, then finally resuspended in FACS buffer and stored at 4°C until analysed.

All data was collected using a Becton Dickinson (BD, Franklin Lakes, NJ, USA) LSR II with standard filter sets and five lasers. Data was analysed using FlowJo software (BD).

Gating strategies are detailed in Supplementary Figure 4.

### List of Flow Antibodies and other reagents

**Table 1.** List of flow cytometry antibodies and reagents.

| Target | Conjugate | Manufacturer | Product Number | Clone | Concentration |
| --- | --- | --- | --- | --- | --- |
| CD45 | PerCP-Cy5.5 | eBioscience | 45-0451 | 30-F11 | 1:400 |
| CD3 | APC | eBioscience | 17-0031 | 145-2C11 | 1:200 |
| CD4 | PE | eBioscience | 12-0041 | GK1.5 | 1:200 |
| CD8 | PE-Cy7 | eBioscience | 561967 | 53-6.7 | 1:400 |
| FoxP3 | FITC | eBioscience | 48-5773 | FJK-165 | 1:100 |
| CD11b | Alexa 700 | eBioscience | 56-0112 | M1/70 | 1:400 |
| F4/80 | PE-Cy5 | eBioscience | 15-4801 | BM8 | 1:400 |
| Ly6G | APC-Cy7 | BD | 560600 | IA8 | 1:200 |
| CD206 | PE | Biolegend | 41705 | C068C2 | 1:100 |
| MHCII | eFluor450 | eBioscience | 48-5321 | M5/114.1.2 | 1:200 |
| Live/Dead | FITC | Invitrogen | L34970 | N/A | 1:200 |

### Fecal DNA Extraction

Feces was weighed into MPBio Lysing Matrix E bead beating tubes (MPBio, Santa Ana, CA, USA) and extraction was completed according to the manufacturer’s protocol for the MPBio FastDNA™ SPIN Kit for Soil but extending the beat beating time to 3 minutes. The DNA recovered from these samples was assessed using a Qubit^®^ 2.0 fluorometer (Invitrogen).

### 16S Library Preparation and Sequencing

Extracted DNA was normalised to 5ng/ul and used in 16S rRNA amplicon PCR targeting the V1+2 of the 16S rRNA gene using the primers detailed in Supplementary Table 3 and the following PCR cycle:

95°C for 3 minutes, 25 cycles of 95°C for 30 seconds, 55°C for 30 seconds and 72°C for 30 seconds with a final 72°C step for 5 minutes. Primer sequences are detailed below.

PCR products were taken through a round of AMPure XP bead clean-up to remove primers and sent to the Wellcome Trust Sanger Institute (Cambridge, UK) for sequencing by Illumina MiSeq 2×300bp paired end chemistry in multiplex generating ~100,000 reads per sample. Raw reads were returned to QIB for analysis.

### Bioinformatic Analysis of 16S sequencing

At QIB, an in-house PE protocol was used for sequencing analysis. After demultiplexing and quality control of raw paired reads using FASTX-Toolkit [52] (minimum quality 33 for at least 50% of the bases in each read sequence) reads were aligned against the SILVA database (version: SILVA_128_SSURef_tax_silva) [53] and BLASTN (ncbi-blast-2.2.25+; Max e-value 10e-3) [54]. BLAST files were imported into MEGAN6 [55] to create proprietary rma6 files using the following parameters: 100 as maximum number of matches per reads, and “Min Score = 50” and “Top Percent = 10”. All rma6 files of paired read sequences were then normalised and compared using MEGAN6.

To make comparisons between study sets, the samples were normalised to the sample with the lowest number of reads. Alpha diversity analyses were performed using MEGAN6. Principal Coordinate Analysis plotting was performed using Bray-Curtis distances from the 16S MEGAN community profiles. Significantly changing genera were identified from relative abundance data using paired and unpaired T-tests with FDR correction at 5%.

### Fecal Metabolomics

Faecal samples were prepared for 1H NMR spectroscopy by mixing 25mg (FW) of faecal samples with 600μL NMR buffer made up of 0.1 M phosphate buffer (0.51 g Na2HPO4, 2.82 g K2HPO4, 100 mg sodium azide and 34.5 mg sodium 3-(Trimethylsilyl)-propionate-d4 (1 mM) in 200 mL deuterium oxide) with a tube pestle. Sample tubes were vortexed for 5 minutes, then centrifuged at 12,000 x g for 5 minutes at 4°C. Supernatant was transferred to a 5-mm NMR tube for recording, conditions were as described below.

Multivariate statistical analyses (Principal Component Analysis) were carried out using the PLS Toolbox v5.5 (Eigenvector Research Inc., Wenatchee, WA) running within Matlab, v7.6 (The MathWorks Inc., Natick, MA) and Metaboanalyst 3.0 [56]. Autoscaling was applied to the columns of the bucket table. Univariate analyses were carried out on individual variates in Excel (t-tests).

### NMR Conditions

High resolution 1H NMR spectra were recorded on a 600MHz Bruker Avance spectrometer fitted with a 5 mm TCI cryoprobe and a 60 slot autosampler (Bruker, Rheinstetten, Germany). Sample temperature was controlled at 300 K. Each spectrum consisted of 1024 scans of 65,536 complex data points with a spectral width of 12.5 ppm (acquisition time 2.67 s). The noesypr1d presaturation sequence was used to suppress the residual water signal with low power selective irradiation at the water frequency during the recycle delay (D1 = 3 s) and mixing time (D8 = 0.01 s). A 90° pulse length of 9.6 μs was set for all samples. Spectra were transformed with 0.3 Hz line broadening and zero filling, manually phased, and baseline corrected using the TOPSPIN 2.0 software. Spectra were transferred into AMIX ^®^ software for bucketing and multivariate analysis applied (using Matlab ^®^ Toolbox software). Spectra were transformed with 0.3 Hz line broadening and zero filling, manually phased, and baseline corrected using the TOPSPIN 2.0 software. Metabolites were identified using information found in the literature (references) or on the web (Human Metabolome Database, http://www.hmdb.ca/) and by use of the 2D-NMR methods, COSY, HSQC, and HMBC.

### Mesoscale Discovery Multiplex Arrays

Tissue samples were weighed into a MPBio Lysing Matrix E bead beating tube (MPBio) with 1ml of homogenisation buffer (PBS + 10% FBS (Invitrogen) + cOmplete™ protease inhibitor (Roche). Tissues were homogenised using an MPBio Fast Prep bead beater at speed 4.0 for 40 seconds followed by speed 6.0 for 40 seconds. Samples were centrifuged at 12,000 x g for 12 minutes at 4°C and subsequently stored at −80°C until analysed. Samples were run on a Mesoscale Discovery (MSD, Rockville, MD, USA) V-PLEX Pro-Inflammatory Panel 1 Mouse Kit according to the manufacturer’s instructions. Plate was read using an MSD QuickPlex SQ 120 imager.

### RNA Sequencing

Whole tumour RNA was extracted as in described previously. Extracted RNA was then quality checked and quantified using a 2100 Bioanalyzer (Agilent) with an RNA 6000 Nano analysis kit (Agilent) and any samples with a RIN value of >8 were considered for use in sequencing. Suitable samples were sent to the Wellcome Trust Sanger Institute for sequencing. All samples were processed by poly-A selection and then sequencing using non-stranded, paired end protocol. Initial processing was performed at Welcome Trust Sanger Institute as follows. Data demultiplexed and adapter removed. Raw reads quality controlled using FastQC (0.11.3, [57]) and trimmed (phred score > 30) using FASTX (0.1.5, [52]). This was followed by read alignment to mouse reference genome (NCBI Mus musculus GRCm38) using Tophat (2.1.1, [58]) using maximum intron size 500.000 bp and default settings. Aligned transcripts were assembled and quantified using Cufflinks (2.1.0, [58]) (applying standard parameters).

At QIB, read alignment and quantification was performed using Kallisto [59]. The quantified read data was then normalised and differential expression analysis was conducted using DeSeq2 [60].. Transcript IDs were annotated using the Ensembl Biomart database. Significantly up and down regulated genes (p_adj_ <0.05) were used to perform biological process and pathway analysis using DAVID. Biological processes were annotated according to the GO_TERM_BP_ALL database and pathway analysis was performed using KEGG pathways. Significantly enriched pathways were determined by a DAVID enrichment score (EASE) of less than 0.05.

### Statistical Analyses

Specific statistical methods are detailed in the relevant methods section or the figure legend. However, unless specified otherwise, statistical analysis was performed using student’s t-test, * p<0.05, ** p<0.01, *** p<0.001.

## Declarations

### Author’s Contributions

LH and SDR conceived of the study, performed experiments, analysed data, interpreted results and participated in writing and review of the manuscript. BMK and AM performed experiments, analysed data, interpreted results and participated in writing and review of the manuscript. KM performed experiments and participated in review of the manuscript. JP performed experiments. CA, CL, MD and SC performed the library preparation and bioinformatic analysis of 16S sequencing. GL performed metabolomic analysis of fecal samples. PK, PD, JM, EC, AG performed corroboration of metabolomic analysis. MY participated in the review of the manuscript. AA and SM provided flow cytometry support, analysed data, and participated in review of the manuscript. KNM provided PyMT-BO1 cells and helped with experimental design. TK provided support with RNA sequencing analysis. All authors read and approved the final manuscript.

## Acknowledgments

We would like to thank Prof. Kairbaan Hodivala-Dilke for supplying the EO771 cells. We would also like to thank Dr Lukas Harnisch for his help with preparing samples for RNA sequencing. This work could not have been completed without the help of everyone in the Robinson and Edwards Labs, including Wesley Fowler, Dr Sally Dreger, Rob Johnson, Jordi Lambert, Abdullah Alghamdi and Aleks Gontarcyzk, plus all previous members who contributed. Additional thanks go to Prof. Dylan Edwards for his guidance (and coffee) throughout.

## Footnotes

a PYMT-BO1 cells obtained from Dr Katherine Weilbaecher (Washington University, St Louis, MO, USA)

b E0771 cells obtained from Prof Kairbaan Hodivala-Dilke (Barts Cancer Institute, QMUL, London, UK)

## References

1. Bray F, Ferlay J, Soerjomataram I, Siegel RL, Torre LA, Jemal A (2018) Global cancer statistics 2018: GLOBOCAN estimates of incidence and mortality worldwide for 36 cancers in 185 countries. CA Cancer J Clin 68:394–424.

2. Allán Pérez-Solis M, Maya-Nuñez G, Casas-González P, Olivares A, Aguilar-Rojas A (2016) Effects of the lifestyle habits in breast cancer transcriptional regulation Cancer Cell International. Cancer Cell Int 16:7.

3. Zitvogel L, Pietrocola F, Kroemer G (2017) Nutrition, inflammation and cancer. Nat Immunol 18:843–850.

4. Kelly D, Mulder IE (2012) Microbiome and immunological interactions. Nutr Rev 70:S18–S30.

5. Hughes ER, Winter MG, Duerkop BA, Spiga L, Furtado de Carvalho T, Zhu W, Gillis CC, Büttner L, Smoot MP, Behrendt CL, Cherry S, Santos RL, Hooper L V., Winter SE (2017) Microbial Respiration and Formate Oxidation as Metabolic Signatures of Inflammation-Associated Dysbiosis. Cell Host Microbe 21:208–219.

6. Zitvogel L, Daillère R, Roberti MP, Routy B, Kroemer G (2017) Anticancer effects of the microbiome and its products. Nat Rev Microbiol 15:465–478.

7. Zitvogel L, Galluzzi L, Viaud S, Vétizou M, Daillère R, Merad M, Kroemer G (2015) Cancer and the gut microbiota: An unexpected link. Sci China Life Sci 7:271ps1.

8. Rooks MG, Garrett WS (2016) Gut microbiota, metabolites and host immunity. Nat Rev Immunol 16:341–352.

9. Wu N, Yang X, Zhang R, et al. (2013) Dysbiosis Signature of Fecal Microbiota in Colorectal Cancer Patients. Microb Ecol 66:462–470.

10. Sivan A, Corrales L, Hubert N, et al. (2015) Commensal Bifidobacterium promotes antitumor immunity and facilitates anti-PD-L1 efficacy.Science 350:1084–9.

11. Aminov RI (2010) A brief history of the antibiotic era: lessons learned and challenges for the future. Front Microbiol 1:134.

12. Becattini S, Taur Y, Pamer EG (2016) Antibiotic-Induced Changes in the Intestinal Microbiota and Disease. Trends Mol Med 22:458–478.

13. Jones DJ, Bunn F, Bell-Syer S V (2014) Prophylactic antibiotics to prevent surgical site infection after breast cancer surgery. Cochrane Database Syst Rev CD005360.

14. Ranganathan K, Sears ED, Zhong L, Chung T-T, Chung KC, Kozlow JH, Momoh AO, Waljee JF (2018) Antibiotic Prophylaxis after Immediate Breast Reconstruction. Plast Reconstr Surg 141:865–877.

15. Tremaroli V, Bäckhed F (2012) Functional interactions between the gut microbiota and host metabolism. Nature 489:242–249.

16. Edwards BL, Stukenborg GJ, Brenin DR, Schroen AT (2014) Use of prophylactic postoperative antibiotics during surgical drain presence following mastectomy. Ann Surg Oncol 21:3249–55.

17. Su X, Esser AK, Amend SR, et al. (2016) Antagonizing Integrin β3 Increases Immunosuppression in Cancer. Cancer Res 76:3484–95.

18. Croswell A, Amir E, Teggatz P, Barman M, Salzman NH (2009) Prolonged impact of antibiotics on intestinal microbial ecology and susceptibility to enteric Salmonella infection. Infect Immun 77:2741–53.

19. Reikvam DH, Erofeev A, Sandvik A, Grcic V, Jahnsen FL, Gaustad P, McCoy KD, Macpherson AJ, Meza-Zepeda LA, Johansen F-E (2011) Depletion of Murine Intestinal Microbiota: Effects on Gut Mucosa and Epithelial Gene Expression. PLoS One 6:e17996.

20. Reivew on Antimicrobial Resistance (2014) Publications | AMR Review. In: Antimicrob. Resist. Tackling a Cris. Futur. Heal. Wealth Nations.https://amr-review.org/Publications.html. Accessed 16 Aug 2018

21. Gopalakrishnan V, Spencer CN, Nezi L, et al. (2018) Gut microbiome modulates response to anti-PD-1 immunotherapy in melanoma patients. Science 359:97–103.

22. Matson V, Fessler J, Bao R, Chongsuwat T, Zha Y, Alegre M-L, Luke JJ, Gajewski TF (2018) The commensal microbiome is associated with anti-PD-1 efficacy in metastatic melanoma patients. Science 359:104–108.

23. Routy B, Le Chatelier E, Derosa L, et al. (2018) Gut microbiome influences efficacy of PD-1-based immunotherapy against epithelial tumors. Science 359:91–97.

24. Vétizou M, Pitt JM, Daillère R, et al. (2015) Anticancer immunotherapy by CTLA-4 blockade relies on the gut microbiota. Science 350:1079–84.

25. Rossini A, Rumio C, Sfondrini L, Tagliabue E, Morelli D, Miceli R, Mariani L, Palazzo M, Ménard S, Balsari A (2006) Influence of antibiotic treatment on breast carcinoma development in proto-neu transgenic mice. Cancer Res 66:6219–24.

26. Naito K, Naito Y, Takagi T, et al. (2011) Role of tumor necrosis factor-α in the pathogenesis of indomethacin-induced small intestinal injury in mice. Int J Mol Med 27:353–9.

27. Lee Y-S, Yang H, Yang J-Y, Kim Y, Lee S-H, Kim JH, Jang YJ, Vallance BA, Kweon M-N (2015) Interleukin-1 (IL-1) Signaling in Intestinal Stromal Cells Controls KC/CXCL1 Secretion, Which Correlates with Recruitment of IL-22-Secreting Neutrophils at Early Stages of Citrobacter rodentium Infection. Infect Immun 83:3257–3267.

28. Wang Y, Cai X, Zhang S, Cui M, Liu F, Sun B, Zhang W, Zhang X, Ye L (2017) HBXIP up-regulates ACSL1 through activating transcriptional factor Sp1 in breast cancer. Biochem Biophys Res Commun 484:565–571.

29. Monaco ME (2017) Fatty acid metabolism in breast cancer subtypes. Oncotarget 8:29487–29500.

30. Chen W-C, Wang C-Y, Hung Y-H, Weng T-Y, Yen M-C, Lai M-D (2016) Systematic Analysis of Gene Expression Alterations and Clinical Outcomes for Long-Chain Acyl-Coenzyme A Synthetase Family in Cancer. PLoS One 11:e0155660.

31. Igal RA (2010) Stearoyl-CoA desaturase-1: a novel key player in the mechanisms of cell proliferation, programmed cell death and transformation to cancer. Carcinogenesis 31:1509–1515.

32. Holder AM, Gonzalez-Angulo AM, Chen H, Akcakanat A, Do K-A, Fraser Symmans W, Pusztai L, Hortobagyi GN, Mills GB, Meric-Bernstam F (2013) High stearoyl-CoA desaturase 1 expression is associated with shorter survival in breast cancer patients. Breast Cancer Res Treat 137:319–327.

33. Luyimbazi D, Akcakanat A, McAuliffe PF, Zhang L, Singh G, Gonzalez-Angulo AM, Chen H, Do K-A, Zheng Y, Hung M-C, Mills GB, Meric-Bernstam F (2010) Rapamycin regulates stearoyl CoA desaturase 1 expression in breast cancer. Mol Cancer Ther 9:2770–84.

34. Chourasia AH, Macleod KF (2015) Tumor suppressor functions of BNIP3 and mitophagy. Autophagy 11:1937–8.

35. Koop EA, van Laar T, van Wichen DF, de Weger RA, van der Wall E, van Diest PJ (2009) Expression of BNIP3 in invasive breast cancer: correlations with the hypoxic response and clinicopathological features. BMC Cancer 9:175.

36. Geerts D, Koster J, Albert D, Koomoa D-LT, Feith DJ, Pegg AE, Volckmann R, Caron H, Versteeg R, Bachmann AS (2010) The polyamine metabolism genes ornithine decarboxylase and antizyme 2 predict aggressive behavior in neuroblastomas with and without MYCN amplification. Int J cancer 126:2012–24.

37. Casero RA, Marton LJ (2007) Targeting polyamine metabolism and function in cancer and other hyperproliferative diseases. Nat Rev Drug Discov 6:373–390.

38. Kovács T, Mikó E, Vida A, Sebő É, Toth J, Csonka T, Boratkó A, Ujlaki G, Lente G, Kovács P, Tóth D, Árkosy P, Kiss B, Méhes G, Goedert JJ, Bai P (2019) Cadaverine, a metabolite of the microbiome, reduces breast cancer aggressiveness through trace amino acid receptors. Sci Rep 9:1300.

39. Davie JR (2003) Inhibition of Histone Deacetylase Activity by Butyrate. J Nutr 133:2485S–2493S.

40. Salimi V, Shahsavari Z, Safizadeh B, Hosseini A, Khademian N, Tavakoli-Yaraki M (2017) Sodium butyrate promotes apoptosis in breast cancer cells through reactive oxygen species (ROS) formation and mitochondrial impairment. Lipids Health Dis 16:208.

41. Waterhouse C, Jeanpretre N, Keilson J (1979) Gluconeogenesis from alanine in patients with progressive malignant disease. Cancer Res 39:1968–72.

42. Kennedy KM, Scarbrough PM, Ribeiro A, Richardson R, Yuan H, Sonveaux P, Landon CD, Chi J-T, Pizzo S, Schroeder T, Dewhirst MW (2013) Catabolism of exogenous lactate reveals it as a legitimate metabolic substrate in breast cancer. PLoS One 8:e75154.

43. Latham T, Mackay L, Sproul D, Karim M, Culley J, Harrison DJ, Hayward L, Langridge-Smith P, Gilbert N, Ramsahoye BH (2012) Lactate, a product of glycolytic metabolism, inhibits histone deacetylase activity and promotes changes in gene expression. Nucleic Acids Res 40:4794–4803.

44. McCafferty J, Mühlbauer M, Gharaibeh RZ, Arthur JC, Perez-Chanona E, Sha W, Jobin C, Fodor AA (2013) Stochastic changes over time and not founder effects drive cage effects in microbial community assembly in a mouse model. ISME J 7:2116–2125.

45. Lim S, Chang D-H, Ahn S, Kim B-C (2016) Whole genome sequencing of “Faecalibaculum rodentium” ALO17, isolated from C57BL/6J laboratory mouse feces. Gut Pathog 8:3.

46. Van den Abbeele P, Belzer C, Goossens M, Kleerebezem M, De Vos WM, Thas O, De Weirdt R, Kerckhof F-M, Van de Wiele T (2013) Butyrate-producing Clostridium cluster XIVa species specifically colonize mucins in an in vitro gut model. ISME J 7:949–61.

47. Vital M, Howe AC, Tiedje JM (2011) Revealing the bacterial butyrate synthesis pathways by analyzing (meta)genomic data. MBio 5:e00889.

48. Lim S, Chang D-H, Ahn S, Kim B-C (2016) Whole genome sequencing of “Faecalibaculum rodentium” ALO17, isolated from C57BL/6J laboratory mouse feces. Gut Pathog 8:3.

49. Bultman SJ, Jobin C (2014) Microbial-derived butyrate: an oncometabolite or tumor-suppressive metabolite? Cell Host Microbe 16:143–145.

50. Edwards R (1997) Resistance to β-lactam antibiotics in bacteroides spp. J Med Microbiol 46:979–986.

51. Chung L, Thiele Orberg E, Geis AL, et al. (2018) Bacteroides fragilis Toxin Coordinates a Pro-carcinogenic Inflammatory Cascade via Targeting of Colonic Epithelial Cells. Cell Host Microbe 23:203–214.e5.

52. Hannon Lab FASTX-Toolkit. http://hannonlab.cshl.edu/fastx_toolkit/index.html. Accessed 28 Aug 2018

53. Quast C, Pruesse E, Yilmaz P, Gerken J, Schweer T, Yarza P, Peplies J, Glöckner FO (2013) The SILVA ribosomal RNA gene database project: improved data processing and web-based tools. Nucleic Acids Res 41:D590–6.

54. Boratyn GM, Camacho C, Cooper PS, Coulouris G, Fong A, Ma N, Madden TL, Matten WT, McGinnis SD, Merezhuk Y, Raytselis Y, Sayers EW, Tao T, Ye J, Zaretskaya I (2013) BLAST: a more efficient report with usability improvements. Nucleic Acids Res 41:W29–W33.

55. Huson DH, Auch AF, Qi J, Schuster SC (2007) MEGAN analysis of metagenomic data. Genome Res 17:377–86.

56. Xia J, Wishart DS (2016) Using MetaboAnalyst 3.0 for Comprehensive Metabolomics Data Analysis. Curr Protoc Bioinforma 55:14.10.1–14.10.91.

57. Andrews S Babraham Bioinformatics - FastQC A Quality Control tool for High Throughput Sequence Data. https://www.bioinformatics.babraham.ac.uk/projects/fastqc/. Accessed 28 Aug 2018

58. Trapnell C, Roberts A, Goff L, Pertea G, Kim D, Kelley DR, Pimentel H, Salzberg SL, Rinn JL, Pachter L (2012) Differential gene and transcript expression analysis of RNA-seq experiments with TopHat and Cufflinks. Nat Protoc 7:562–78.

59. Bray NL, Pimentel H, Melsted P, Pachter L (2016) Near-optimal probabilistic RNA-seq quantification. Nat Biotechnol 34:525–527.

60. Love MI, Huber W, Anders S (2014) Moderated estimation of fold change and dispersion for RNA-seq data with DESeq2. Genome Biol 15:550.

