## Supplementary Figures and Tables for "Perturbation of the gut microbiota by antibiotics results in accelerated breast tumour growth and metabolic dysregulation"

**Supplementary Figure 1 – Effect of VNMA administration on intestinal cytokine production:** Cytokines listed were analysed by MSD V-Plex assay and normalised to tissue weight. Data displayed in log scale to account for large differences in concentration between cytokines. Significance was determined by unpaired, two-sample T test, * $p<0.05$, ** $p<0.01$, $n \geq 5$ per condition.

**Supplementary Figure 2 – Full list of differentially expressed genes from RNAseq analysis:** A) Genes significantly downregulated in VNMA treated tumours. B) Genes significantly upregulated in VNMA treated tumours

**Supplementary Figure 3 – Heatmaps showing DEGS involved in other biological functions:** Gene clusters identified by RNAseq analysis, grouped by their overarching biological function according to GO classifiers.

**Supplementary Figure 4 – Gating strategies used in flow cytometry:** Pathway for identification of immune cells, cell debris, doublets and dead cells are first excluded. Leukocytes are selected as $CD45^{+}$ events before further delineation to either myeloid cells (A) or T cells (B). Gating shown in B occurs after the previously mentioned gating steps.

**Supplementary Figure 5 – Heatmap of full list of fecal metabolites identified by NMR:** Sample A3 in the control group was shown to be a significant outlier across several metabolites and was therefore excluded from further analysis.

Supplementary Figure 1

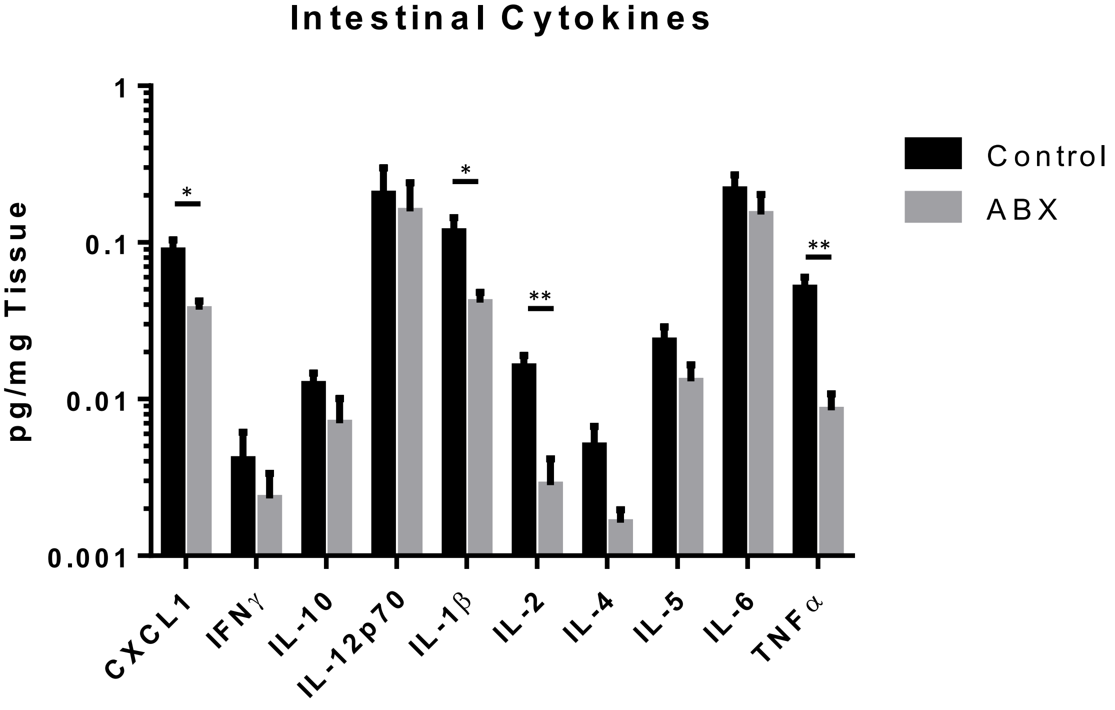

### Supplementary Figure 2

**A**

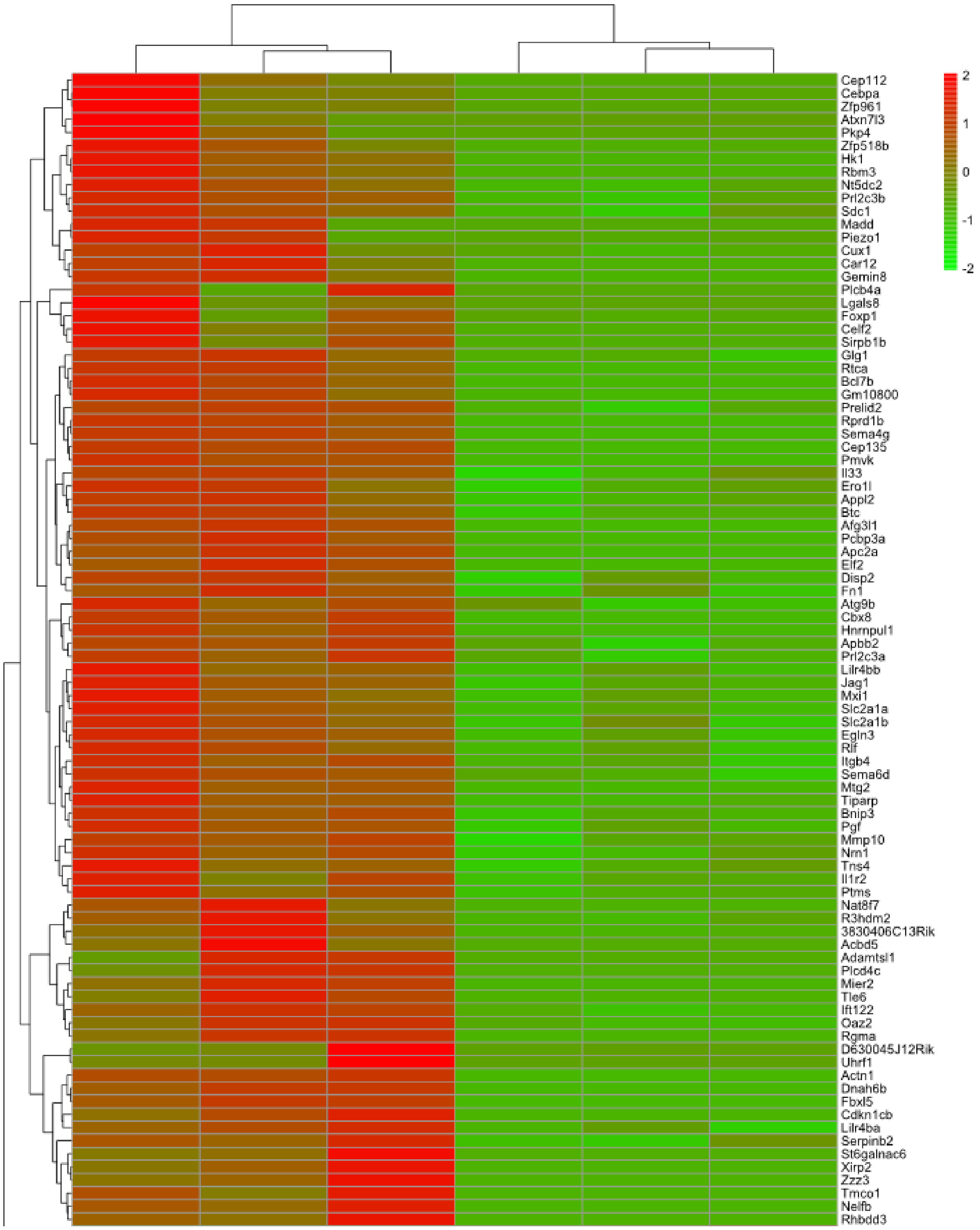

**B**

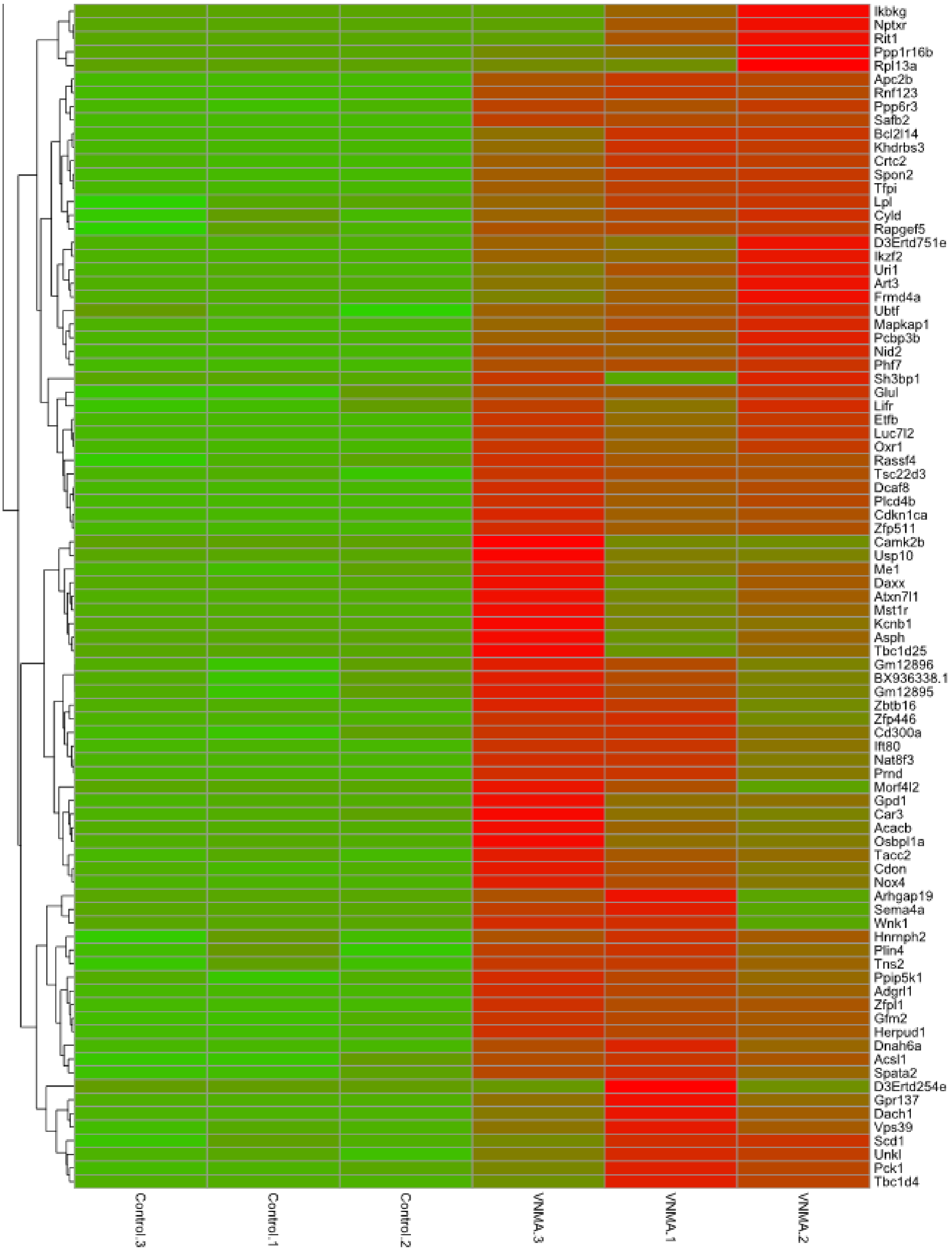
Ikbkg
Nptxr
Rit1
Ppp1r16b
Rpl13a
Apc2b
Rnf123
Ppp6r3
Safb2
Bcl2l14
Khdrbs3
Crtc2
Spon2
Tfpi
Lpl
Cyld
Rapgef5
D3Ertd751e
Ikzf2
Uri1
Art3
Frmd4a
Ubtf
Mapkap1
Pcbp3b
Nid2
Phf7
Sh3bp1
Glul
Lifr
Etfb
Luc7l2
Oxr1
Rassf4
Tsc22d3
Dcaf8
Plcd4b
Cdkn1ca
Zfp511
Camk2b
Usp10
Me1
Daxx
Atxn7l1
Mst1r
Kcnb1
Asph
Tbc1d25
Gm12896
BX936338.1
Gm12895
Zbtb16
Zfp446
Cd300a
Ift80
Nat8f3
Prnd
Morf4l2
Gpd1
Car3
Acacb
Osbpl1a
Tacc2
Cdon
Nox4
Arhgap19
Sema4a
Wnk1
Hnrnph2
Plin4
Tns2
Ppip5k1
Adgrl1
Zfpl1
Gfm2
Herpud1
Dnah6a
Acsl1
Spata2
D3Ertd254e
Gpr137
Dach1
Vps39
Scd1
Unkl
Pck1
Tbc1d4
Control.3
Control.1
Control.2
VNMA.3
VNMA.1
VNMA.2

### Supplementary Figure 3

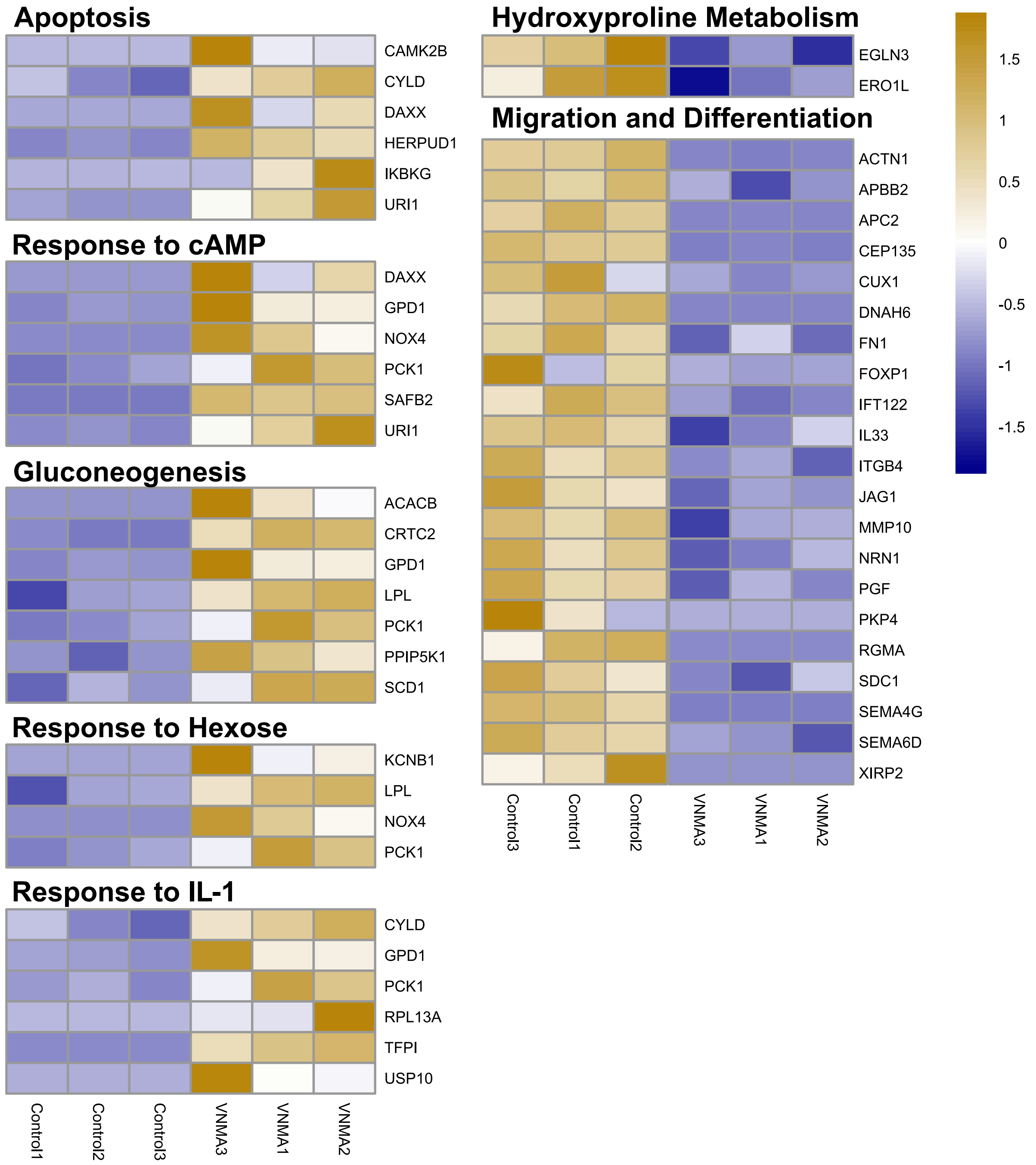

Supplementary Figure 4A

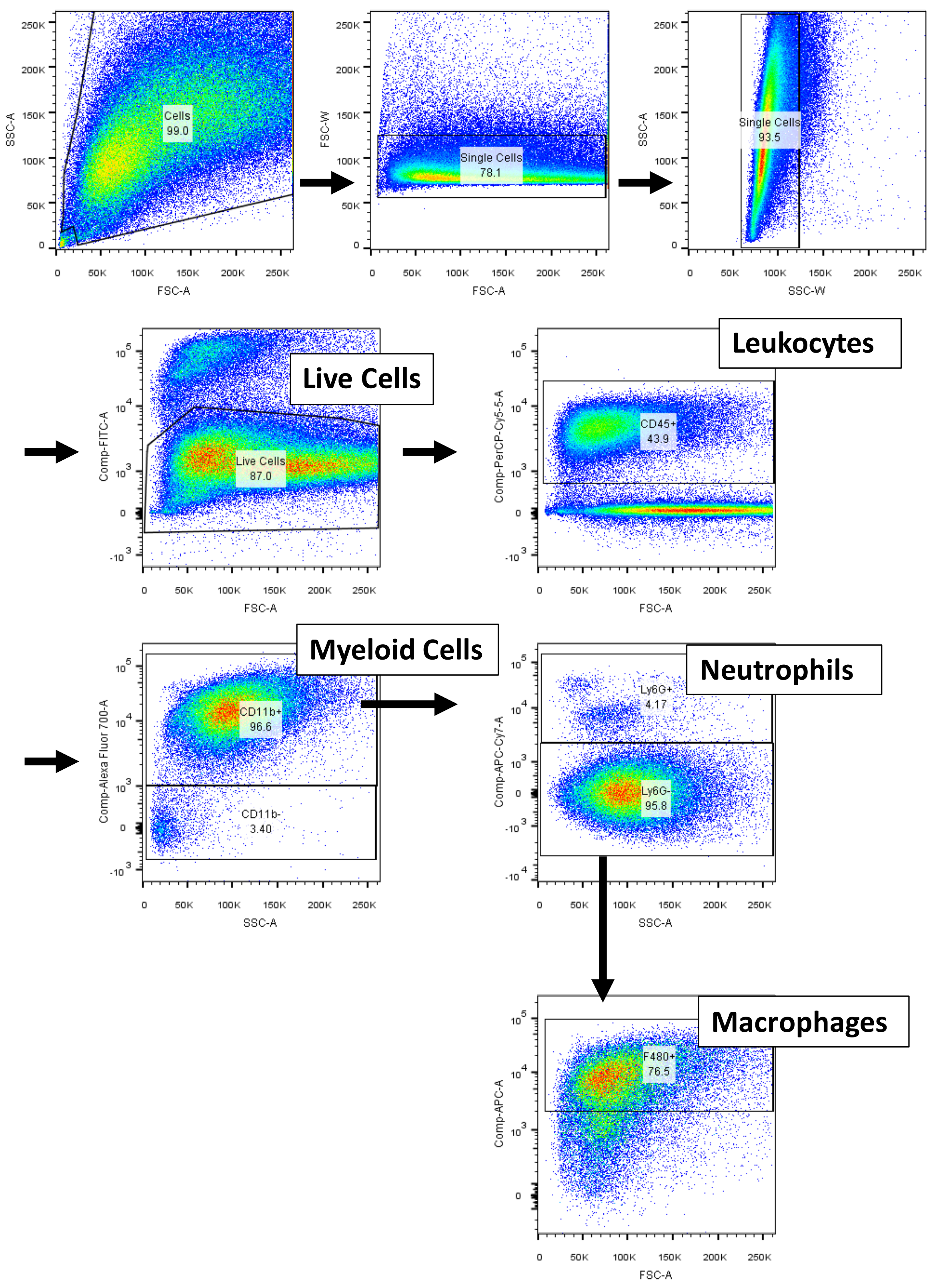

Supplementary Figure 4B

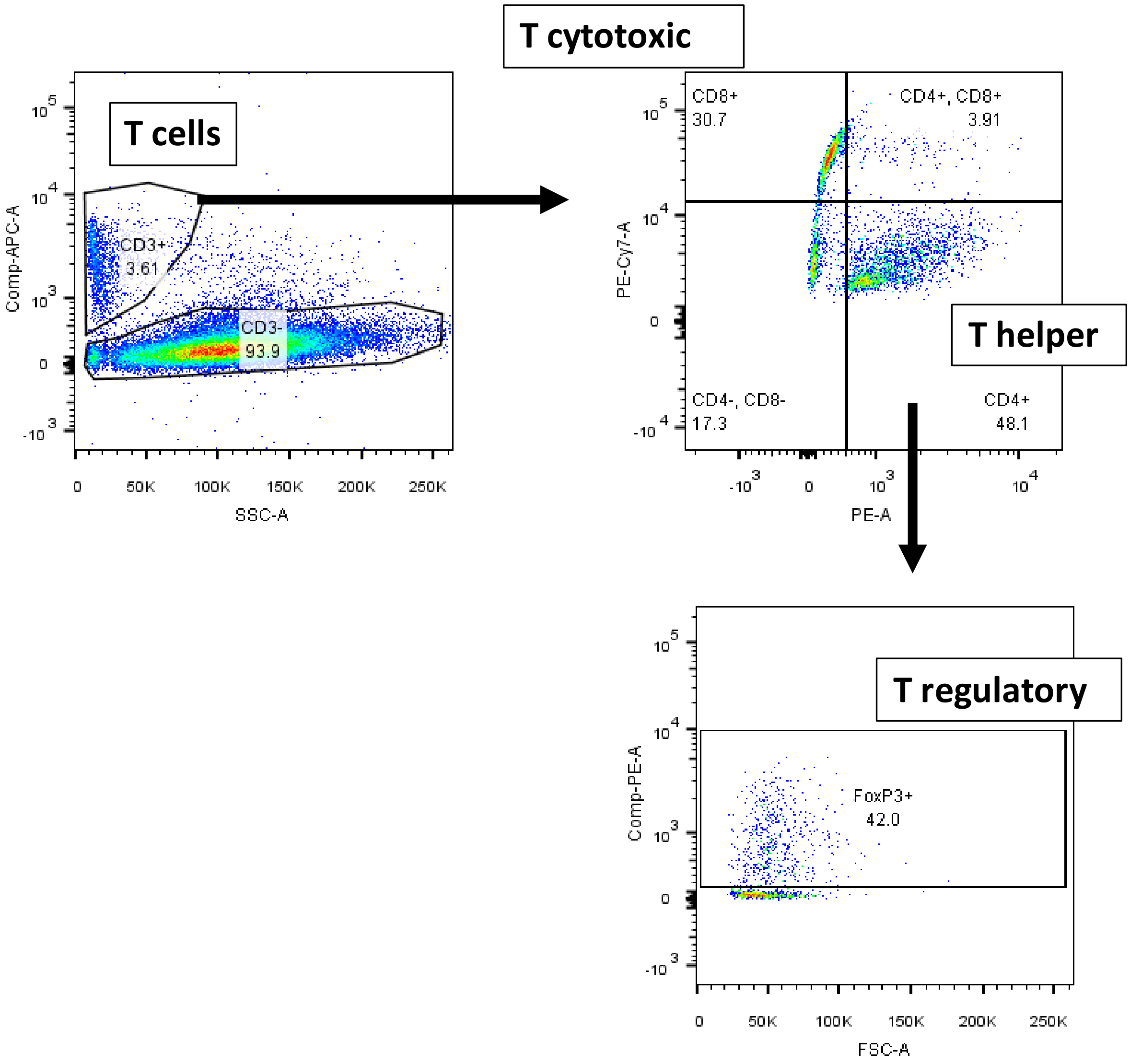

### Supplementary Figure 5

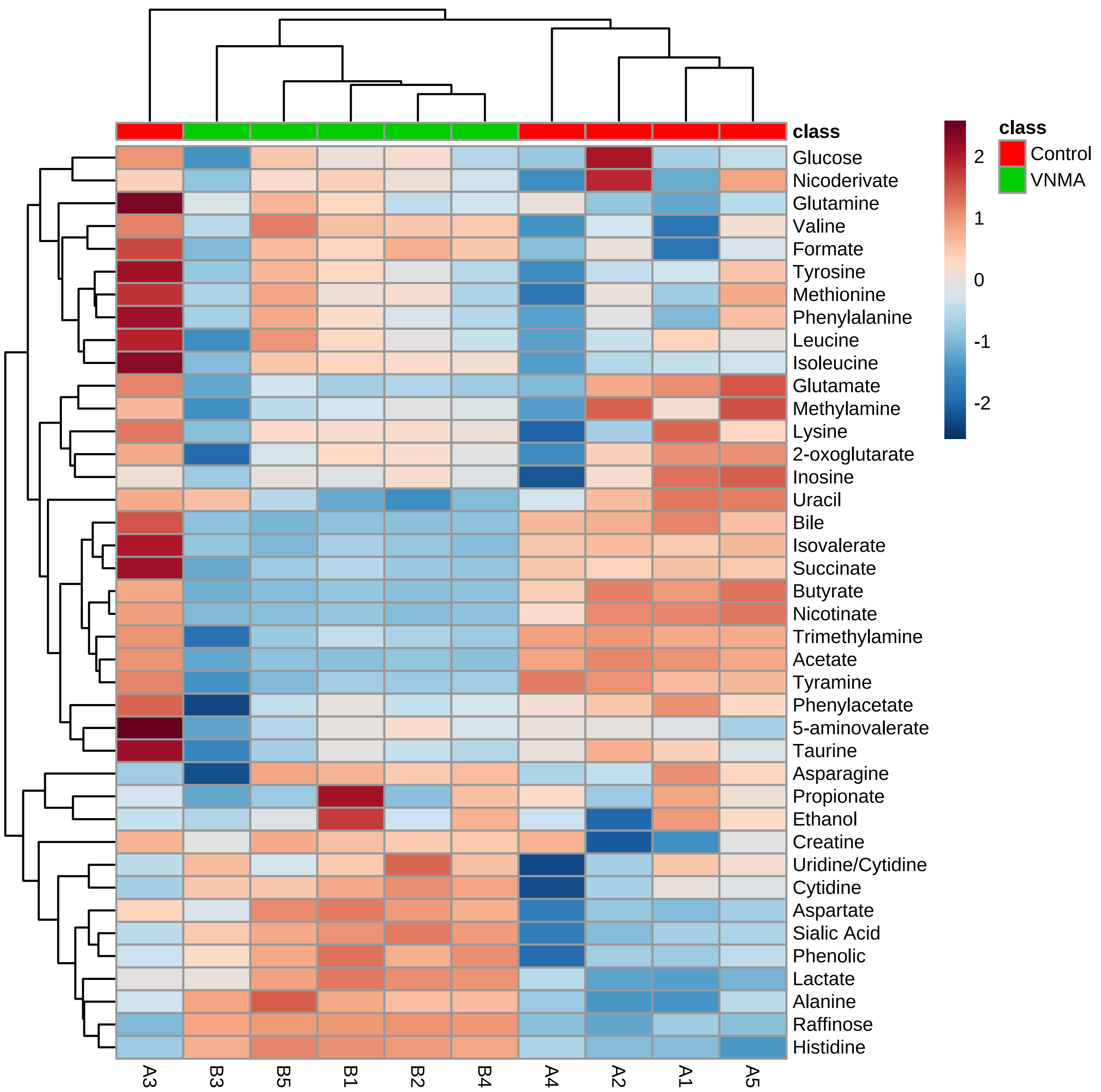

#### Table 1 – Full list of biological processes upregulated in VNMA tumours

| Term | Count | PValue |
|---|---|---|
| GO:0071704~organic substance metabolic process | 52 | 0.044923318 |
| GO:0044237~cellular metabolic process | 51 | 0.018768574 |
| GO:0044238~primary metabolic process | 51 | 0.021510919 |
| GO:0051716~cellular response to stimulus | 39 | 0.040996104 |
| GO:0019222~regulation of metabolic process | 36 | 0.013468412 |
| GO:0080090~regulation of primary metabolic process | 35 | 0.006703144 |
| GO:0031323~regulation of cellular metabolic process | 35 | 0.007640595 |
| GO:0016043~cellular component organization | 34 | 0.023968801 |
| GO:0071840~cellular component organization or biogenesis | 34 | 0.035533607 |
| GO:0048519~negative regulation of biological process | 33 | 0.001525579 |
| GO:0048518~positive regulation of biological process | 33 | 0.009594897 |
| GO:0048523~negative regulation of cellular process | 32 | 0.000911725 |
| GO:0048522~positive regulation of cellular process | 30 | 0.014753971 |
| GO:0044267~cellular protein metabolic process | 27 | 0.048181807 |
| GO:0042221~response to chemical | 26 | 0.022217187 |
| GO:0010033~response to organic substance | 23 | 0.003690949 |
| GO:0036211~protein modification process | 23 | 0.022564955 |
| GO:0006464~cellular protein modification process | 23 | 0.022564955 |
| GO:0043412~macromolecule modification | 23 | 0.045212693 |
| GO:0009893~positive regulation of metabolic process | 21 | 0.018424214 |
| GO:0031325~positive regulation of cellular metabolic process | 20 | 0.016243506 |
| GO:0065009~regulation of molecular function | 19 | 0.003288413 |
| GO:0010646~regulation of cell communication | 19 | 0.033561161 |
| GO:0010604~positive regulation of macromolecule metabolic process | 19 | 0.034524527 |
| GO:0023051~regulation of signaling | 19 | 0.036064428 |
| GO:0070887~cellular response to chemical stimulus | 18 | 0.036148898 |
| GO:0071310~cellular response to organic substance | 17 | 0.011353823 |
| GO:0043933~macromolecular complex subunit organization | 17 | 0.019021129 |
| GO:0031324~negative regulation of cellular metabolic process | 17 | 0.03295039 |
| GO:0035556~intracellular signal transduction | 17 | 0.03295039 |
| GO:0009966~regulation of signal transduction | 17 | 0.043995234 |
| GO:0050790~regulation of catalytic activity | 16 | 0.003051297 |
| GO:0042592~homeostatic process | 15 | 0.009773798 |
| GO:0019220~regulation of phosphate metabolic process | 14 | 0.016054466 |
| GO:0051174~regulation of phosphorus metabolic process | 14 | 0.016129513 |
| GO:1901700~response to oxygen-containing compound | 14 | 0.019101669 |
| GO:0006357~regulation of transcription from RNA polymerase II promoter | 14 | 0.029706337 |
| GO:0008283~cell proliferation | 14 | 0.043516668 |
| GO:0044093~positive regulation of molecular function | 13 | 0.005795768 |
| GO:0042981~regulation of apoptotic process | 13 | 0.018395437 |
| GO:0043067~regulation of programmed cell death | 13 | 0.019849603 |
| GO:0010941~regulation of cell death | 13 | 0.033948685 |
| GO:0045935~positive regulation of nucleobase-containing compound metabolic process | 13 | 0.036376781 |
| GO:0006366~transcription from RNA polymerase II promoter | 13 | 0.046468607 |
| GO:0048585~negative regulation of response to stimulus | 12 | 0.022886977 |
| GO:0010648~negative regulation of cell communication | 11 | 0.021101866 |
| GO:0023057~negative regulation of signaling | 11 | 0.021556067 |
| GO:0006082~organic acid metabolic process | 10 | 0.010245582 |
| GO:0051336~regulation of hydrolase activity | 10 | 0.011597597 |
| GO:0043085~positive regulation of catalytic activity | 10 | 0.01805321 |
| GO:0009968~negative regulation of signal transduction | 10 | 0.025540497 |
| GO:0019752~carboxylic acid metabolic process | 9 | 0.016504911 |
| GO:0043436~oxoacid metabolic process | 9 | 0.017195776 |
| GO:0044283~small molecule biosynthetic process | 8 | 0.002775214 |
| GO:0071345~cellular response to cytokine stimulus | 8 | 0.007157341 |
| GO:0008285~negative regulation of cell proliferation | 8 | 0.017929439 |
| GO:0034097~response to cytokine | 8 | 0.021559377 |
| GO:1902532~negative regulation of intracellular signal transduction | 7 | 0.011938911 |
| GO:0032787~monocarboxylic acid metabolic process | 7 | 0.027131369 |
| GO:0046486~glycerolipid metabolic process | 6 | 0.006895312 |
| GO:2001233~regulation of apoptotic signaling pathway | 6 | 0.021197503 |
| GO:0045860~positive regulation of protein kinase activity | 6 | 0.022494599 |
| GO:0033674~positive regulation of kinase activity | 6 | 0.031092398 |
| GO:0035303~regulation of dephosphorylation | 5 | 0.004122138 |
| GO:0009746~response to hexose | 5 | 0.011492944 |
| GO:0034284~response to monosaccharide | 5 | 0.012631472 |
| GO:0009743~response to carbohydrate | 5 | 0.017414907 |
| GO:0051260~protein homooligomerization | 5 | 0.035679943 |
| GO:0006641~triglyceride metabolic process | 4 | 0.006808079 |
| GO:0006639~acylglycerol metabolic process | 4 | 0.010770246 |
| GO:0006638~neutral lipid metabolic process | 4 | 0.011307438 |
| GO:0010921~regulation of phosphatase activity | 4 | 0.01360947 |
| GO:2001242~regulation of intrinsic apoptotic signaling pathway | 4 | 0.024051754 |
| GO:0006475~internal protein amino acid acetylation | 4 | 0.0244711 |
| GO:0006473~protein acetylation | 4 | 0.03834565 |
| GO:0071320~cellular response to cAMP | 3 | 0.01828809 |
| GO:0006094~gluconeogenesis | 3 | 0.035092061 |
| GO:0019319~hexose biosynthetic process | 3 | 0.037841663 |
| GO:0046364~monosaccharide biosynthetic process | 3 | 0.039719255 |
| GO:0035304~regulation of protein dephosphorylation | 3 | 0.043578163 |
| GO:0035305~negative regulation of dephosphorylation | 3 | 0.048589269 |
| GO:0043648~dicarboxylic acid metabolic process | 3 | 0.049615651 |
| GO:0090129~positive regulation of synapse maturation | 2 | 0.044703048 |

#### Table 2 – Full list of biological processes downregulated in VNMA tumours

| Term | Count | PValue |
|---|---|---|
| GO:0043170~macromolecule metabolic process | 47 | 0.030204 |
| GO:0044260~cellular macromolecule metabolic process | 43 | 0.037549 |
| GO:0019222~regulation of metabolic process | 37 | 0.009313 |
| GO:0080090~regulation of primary metabolic process | 36 | 0.004409 |
| GO:0060255~regulation of macromolecule metabolic process | 36 | 0.006201 |
| GO:0048518~positive regulation of biological process | 35 | 0.003153 |
| GO:0016043~cellular component organization | 35 | 0.016832 |
| GO:0071840~cellular component organization or biogenesis | 35 | 0.025598 |
| GO:0010467~gene expression | 33 | 0.0093 |
| GO:0051179~localization | 33 | 0.027533 |
| GO:0031323~regulation of cellular metabolic process | 33 | 0.03111 |
| GO:0048519~negative regulation of biological process | 32 | 0.004131 |
| GO:0048522~positive regulation of cellular process | 31 | 0.009611 |
| GO:0048523~negative regulation of cellular process | 30 | 0.005347 |
| GO:0010468~regulation of gene expression | 28 | 0.012674 |
| GO:0048869~cellular developmental process | 28 | 0.022581 |
| GO:0030154~cell differentiation | 26 | 0.029806 |
| GO:2000112~regulation of cellular macromolecule biosynthetic process | 24 | 0.044875 |
| GO:0010604~positive regulation of macromolecule metabolic process | 21 | 0.009994 |
| GO:0009893~positive regulation of metabolic process | 21 | 0.021209 |
| GO:0009653~anatomical structure morphogenesis | 20 | 0.017048 |
| GO:0007166~cell surface receptor signaling pathway | 18 | 0.012319 |
| GO:0009888~tissue development | 15 | 0.024624 |
| GO:0006928~movement of cell or subcellular component | 14 | 0.026699 |
| GO:0006357~regulation of transcription from RNA polymerase II promoter | 14 | 0.032735 |
| GO:0008283~cell proliferation | 14 | 0.047717 |
| GO:0010629~negative regulation of gene expression | 13 | 0.038373 |
| GO:0051674~localization of cell | 12 | 0.024779 |
| GO:0048870~cell motility | 12 | 0.024779 |
| GO:0051247~positive regulation of protein metabolic process | 12 | 0.035757 |
| GO:0000902~cell morphogenesis | 11 | 0.041743 |
| GO:0048858~cell projection morphogenesis | 10 | 0.007869 |
| GO:0032990~cell part morphogenesis | 10 | 0.010022 |
| GO:0035295~tube development | 9 | 0.009158 |
| GO:0000122~negative regulation of transcription from RNA polymerase II promoter | 9 | 0.013278 |
| GO:0032101~regulation of response to external stimulus | 9 | 0.015239 |
| GO:0030163~protein catabolic process | 9 | 0.015457 |
| GO:0000904~cell morphogenesis involved in differentiation | 9 | 0.019263 |
| GO:0048667~cell morphogenesis involved in neuron differentiation | 8 | 0.006764 |
| GO:0048812~neuron projection morphogenesis | 8 | 0.009985 |
| GO:0045732~positive regulation of protein catabolic process | 7 | 0.000107 |
| GO:0009896~positive regulation of catabolic process | 7 | 0.000771 |
| GO:0042176~regulation of protein catabolic process | 7 | 0.001997 |
| GO:0072001~renal system development | 7 | 0.002267 |
| GO:0001655~urogenital system development | 7 | 0.004274 |
| GO:0007409~axonogenesis | 7 | 0.008534 |
| GO:0061564~axon development | 7 | 0.012199 |
| GO:0009894~regulation of catabolic process | 7 | 0.012693 |
| GO:0051051~negative regulation of transport | 7 | 0.020285 |
| GO:0001822~kidney development | 6 | 0.008727 |
| GO:0016049~cell growth | 6 | 0.04633 |
| GO:0034330~cell junction organization | 5 | 0.009859 |
| GO:0048588~developmental cell growth | 5 | 0.012807 |
| GO:0051224~negative regulation of protein transport | 5 | 0.015129 |
| GO:0060560~developmental growth involved in morphogenesis | 5 | 0.016267 |
| GO:1904950~negative regulation of establishment of protein localization | 5 | 0.017702 |
| GO:0051961~negative regulation of nervous system development | 5 | 0.045789 |
| GO:0014032~neural crest cell development | 4 | 0.00433 |
| GO:0014033~neural crest cell differentiation | 4 | 0.005505 |
| GO:0061387~regulation of extent of cell growth | 4 | 0.013179 |
| GO:0050709~negative regulation of protein secretion | 4 | 0.019634 |
| GO:1990138~neuron projection extension | 4 | 0.028471 |
| GO:0034329~cell junction assembly | 4 | 0.029879 |
| GO:0014031~mesenchymal cell development | 4 | 0.036402 |
| GO:0050770~regulation of axonogenesis | 4 | 0.03909 |
| GO:0050663~cytokine secretion | 4 | 0.03909 |
| GO:0030308~negative regulation of cell growth | 4 | 0.040751 |
| GO:0048762~mesenchymal cell differentiation | 4 | 0.043597 |
| GO:0008361~regulation of cell size | 4 | 0.048353 |
| GO:0050873~brown fat cell differentiation | 3 | 0.011114 |
| GO:0001755~neural crest cell migration | 3 | 0.023972 |
| GO:0050771~negative regulation of axonogenesis | 3 | 0.026373 |
| GO:0007044~cell-substrate junction assembly | 3 | 0.039719 |
| GO:0051781~positive regulation of cell division | 3 | 0.04264 |
| GO:0019471~4-hydroxyproline metabolic process | 2 | 0.041255 |

**Table 3 – Primers used for 16S sequencing**

| Primer Name | Sequence |
|---|---|
| **V1FW_SD501** | AATGATACGGCGACCACCGAGATCTACACAAGCAGCATATGGTAATTGTAGMGTTYGATYMTGGCTCAG |
| **V1FW_SD502** | AATGATACGGCGACCACCGAGATCTACACACGCGTGATATGGTAATTGTAGMGTTYGATYMTGGCTCAG |
| **V1FW_SD503** | AATGATACGGCGACCACCGAGATCTACACCGATCTACTATGGTAATTGTAGMGTTYGATYMTGGCTCAG |
| **V1FW_SD504** | AATGATACGGCGACCACCGAGATCTACACTGCGTCACTATGGTAATTGTAGMGTTYGATYMTGGCTCAG |
| **V1FW_SD505** | AATGATACGGCGACCACCGAGATCTACACGTCTAGTGTATGGTAATTGTAGMGTTYGATYMTGGCTCAG |
| **V1FW_SD506** | AATGATACGGCGACCACCGAGATCTACACCTAGTATGTATGGTAATTGTAGMGTTYGATYMTGGCTCAG |
| **V1FW_SD507** | AATGATACGGCGACCACCGAGATCTACACGATAGCGTTATGGTAATTGTAGMGTTYGATYMTGGCTCAG |
| **V1FW_SD508** | AATGATACGGCGACCACCGAGATCTACACTCTACACTTATGGTAATTGTAGMGTTYGATYMTGGCTCAG |
| **V1FW_SA501** | AATGATACGGCGACCACCGAGATCTACACATCGTACGTATGGTAATTGTAGMGTTYGATYMTGGCTCAG |
| **V2RV_SD701** | CAAGCAGAAGACGGCATACGAGATACCTAGTAAGTCAGTCAGCCGCTGCCTCCCGTAGGAGT |
| **V2RV_SD702** | CAAGCAGAAGACGGCATACGAGATACGTACGTAGTCAGTCAGCCGCTGCCTCCCGTAGGAGT |
| **V2RV_SD703** | CAAGCAGAAGACGGCATACGAGATATATCGCGAGTCAGTCAGCCGCTGCCTCCCGTAGGAGT |
| **V2RV_SD704** | CAAGCAGAAGACGGCATACGAGATCACGATAGAGTCAGTCAGCCGCTGCCTCCCGTAGGAGT |
| **V2RV_SD705** | CAAGCAGAAGACGGCATACGAGATCGTATCGCAGTCAGTCAGCCGCTGCCTCCCGTAGGAGT |
| **V2RV_SD706** | CAAGCAGAAGACGGCATACGAGATCTGCGACTAGTCAGTCAGCCGCTGCCTCCCGTAGGAGT |
| **V2RV_SD707** | CAAGCAGAAGACGGCATACGAGATGCTGTAACAGTCAGTCAGCCGCTGCCTCCCGTAGGAGT |
| **V2RV_SD708** | CAAGCAGAAGACGGCATACGAGATGGACGTTAAGTCAGTCAGCCGCTGCCTCCCGTAGGAGT |
| **V2RV_SD709** | CAAGCAGAAGACGGCATACGAGATGGTCGTAGAGTCAGTCAGCCGCTGCCTCCCGTAGGAGT |
| **V2RV_SD710** | CAAGCAGAAGACGGCATACGAGATTAAGTCTCAGTCAGTCAGCCGCTGCCTCCCGTAGGAGT |
| **V2RV_SD711** | CAAGCAGAAGACGGCATACGAGATTACACAGTAGTCAGTCAGCCGCTGCCTCCCGTAGGAGT |
| **V2RV_SD712** | CAAGCAGAAGACGGCATACGAGATTTGACGCAAGTCAGTCAGCCGCTGCCTCCCGTAGGAGT |
